# HP1-enhanced chromatin compaction stabilizes a synthetic metabolic circuit in yeast

**DOI:** 10.1101/2025.03.04.641524

**Authors:** Lidice González, Sergio A. García Echauri, Célia Jeronimo, Christian Poitras, Melis Gencel, Adrian Serohijos, Kerry Bloom, François Robert, José L. Avalos, Stephen W. Michnick

## Abstract

Chromatin compaction defines genome topology, evolution, and function. The Saccharomycotina subphylum, including the fermenting yeast *Saccharomyces cerevisiae* have a decompacted genome, possibly because they lost two genes mediating a specific histone lysine methylation and histone binding protein heterochromatin protein 1 (HP1). This decompaction may result in the higher-than-expected mutation and meiotic recombination rates observed in this species. To test this hypothesis, we retro-engineered *S. cerevisiae* to compact the genome by expressing the HP1 homologue of *Schizosaccharomyces pombe Sp*Swi6 and H3K9 methyltransferase *Sp*Clr4. The resulting strain had significantly more compact chromatin and reduced rates of mutation and meiotic recombination. The increased genomic stability significantly prolongs the optogenetic control of an engineered strain designed to grow only in blue light. This result demonstrates the potential of our approach to enhance the stability of strains for metabolic engineering and other synthetic biology applications, which are prone to lose activities due to genetic instability.

## Introduction

The recent reports of the substitution of half of the yeast *Saccharomyces cerevisiae* (*S. cerevisiae*) chromosomes with completely synthetic and edited ones, marks a milestone in the field of synthetic biology and promises possibilities of new insights into chromatin biology^1–3^. Among the main goals of this effort is to eliminate potential sources of instability in the yeast genome that cause disruption of metabolically engineered strains by removing genomic elements susceptible to mutation. DNA compaction is essential to control genome organization, expression, and stability to mutation. Well known examples of chromatin organization comprise the existence of chromosome territories as well as the separation into domains of transcriptionally active and repressed regions, including the compartmentalization of the ribosomal DNA genes^4^ and other structural elements such as the topologically associating domains (TADs)^5–7^. These domains, found in many organisms and ranging from hundreds of kilobases to several megabases in length, are believed to constitute regulatory regions defined by specific contacts between different DNA elements to form enhancers, promoters, and heterochromatin components among others^5,8–11^.

The formation of heterochromatin domains regulates genome organization in different ways, for instance, by modulating gene expression and changing the physicochemical properties of the nucleoplasm^12,13^. A well characterized and abundant type of heterochromatin in eukaryotes contains the heterochromatin protein 1 (HP1), a chromodomain-containing protein that binds histone H3 methylated on lysine 9 (H3K9me) among other components^14–17^. This mechanism of heterochromatin formation is well conserved in most eukaryotic organisms^18^. Furthermore, it was recently demonstrated that the *Schizosaccharomyces pombe* (*S. pombe*) homologue of HP1, *Sp*Swi6 induces self-association and compaction of nucleosome arrays and undergoes phase separation to form high density chromatin biomolecular condensates^17^.

Fungi of the Ascomycota subphylum Saccharomycotina lack the lysine 9-methylated histone H3 form of eukaryotic heterochromatin, resulting in mostly decompacted chromatin^19,20^ because they lost the ancestral genes encoding the present-day *S. pombe Sp*Clr4 (H3K9 methyltransferase), and *Sp*Swi6 (HP1) about 300 million years ago (Fig. 1a). *S. cerevisiae* exhibits less heterochromatic regions than *S. pombe*^21,22^. Interestingly, *S. pombe* is considered to have chromosome condensation similar to the one in human cells^21,23,24^. Studies haven shown that *S. cerevisiae* does not form compact structures in interphase or mitosis, consistent with the fact that most of its genes are in an open configuration and either actively transcribed or primed to be rapidly induced^22,25^. Initial Hi-C experiments in this organism did not reveal TAD structures^7^, while abundant 50-100 Kb TADs have been found when analyzing *S. pombe* genome^26^. Although more recent results from *S. cerevisiae* obtained with Micro-C techniques have shown the presence of self-associating regions, these are much shorter with an approximate size of 5 kb^27^. Additionally, the presence of heterochromatin in budding yeast is mainly constrained to the mating type loci, telomeric regions, and the ribosomal DNA array^19^.

**Fig. 1.**
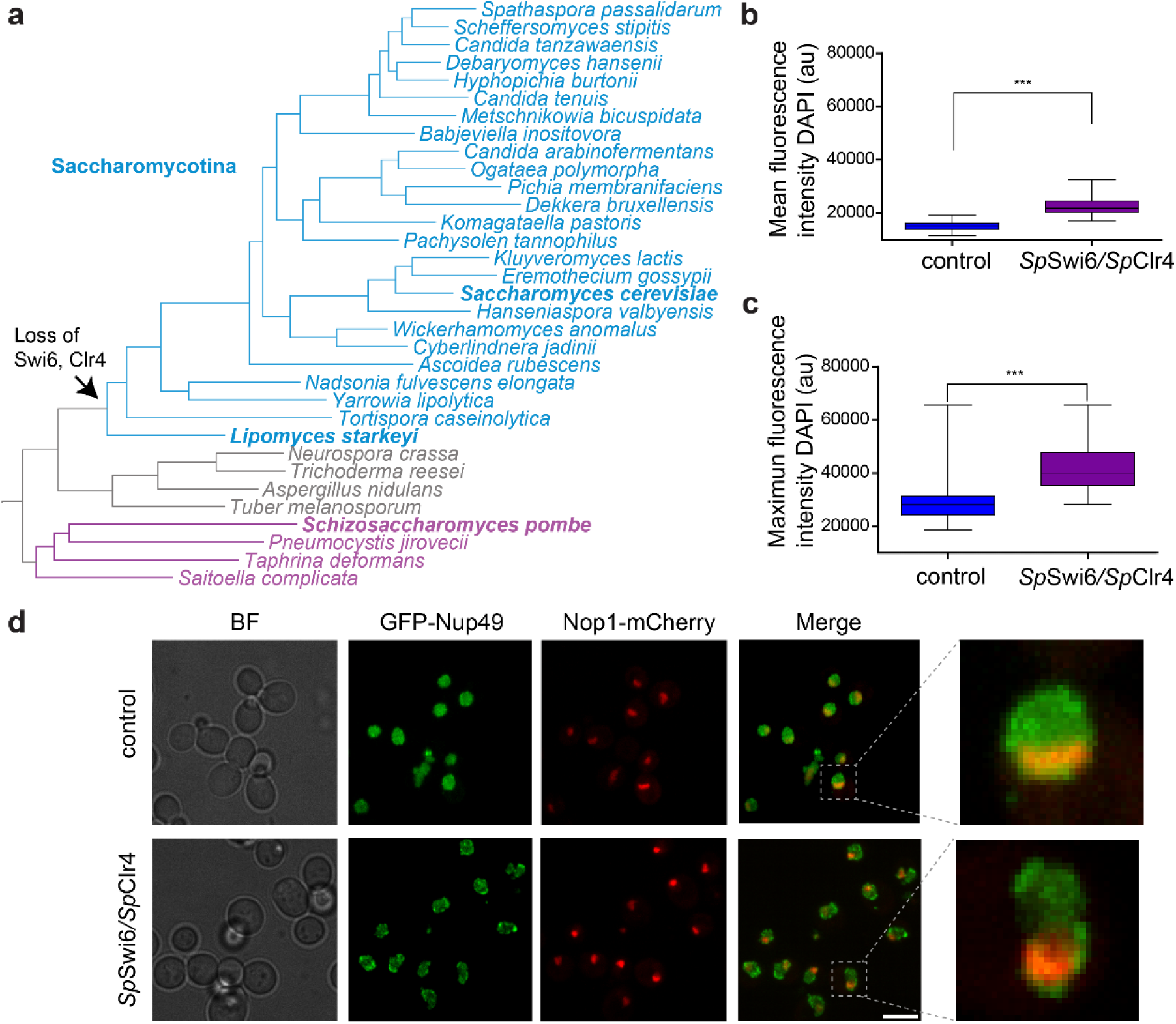
Expressing SpSwi6/SpClr4 proteins increases chromatin compaction in S. cerevisiae. **a,** Phylogenetic tree of yeast species. The arrow indicates where yeast HP1 and H3K9 methyltransferases were lost during evolution, ∼300 million years ago. Blue and purple highlighted regions indicate organisms mentioned in this study (modified from Riley et al., 2016^37^). **b**, **c**, Mean and maximum fluorescence intensity of chromatin stained with DAPI significantly increases in SpSwi6/SpClr4 expressing cells versus reference W303 (two-sided t-test, p < 0.0001). **d**, Live cell imaging of GFP-Nup49 (green) and Nop1-mCherry (red) nuclear envelope (NE) and nucleolus (NO) markers, respectively, in reference (TMS1-1A strain containing plasmids pASZ11-NupNop, pCM189, and p413) and cells expressing the SpSwi6/SpClr4 proteins (TMS1-1A strain containing plasmids pASZ11-NupNop, pCM189-SpSwi6, and p413-SpClr4) (maximum projection of 250 nm Z stacks). BF, brightfield images of the cells. Grey dashed square highlights representative nuclei in both reference and SpSwi6/SpClr4 expressing cells. In b-c, the median is indicated as middle line, 25th and 75th percentile as boxes and the whiskers minimum and maximum values. Scale bar represents 5 µm.

We hypothesized that the *S. cerevisiae* chromatin would become more compact and stable to mutation by expressing *Spswi6* and *Spclr4* (*Spswi6/Spclr4*). We demonstrate that expressing *Sp*Swi6/*Sp*Clr4 does increase compaction of chromatin and reduces both mutation and recombination rates. We further show that strains expressing *Sp*Swi6/*Sp*Clr4 can significantly extend the stability of engineered optogenetic controls of cell growth.

## Results

### *Sp*Swi6/*Sp*Clr4 expression induces global chromatin compaction

Three observations suggest that *Sp*Swi6/*Sp*Clr4 expression causes compaction of yeast chromatin. *Spswi6* and *Spclr4* were expressed in *S. cerevisiae* under control of tunable promoters TetO (7 sites) and *MET17*, respectively. First, we measured chromatin compaction by quantifying 4′,6-diamidino-2-phenylindole (DAPI) binding to the minor groove of DNA, which is sensitive to chromatin structure and increases with chromatin compaction^28,29^. For example, highly condensed pericentric heterochromatin has been shown to form dense foci easily detectable by DNA staining^30–32^. These regions appear also marked with HP1^30,33^, the associated H3K9me^34^, as well as methyl-CpG binding protein 2 (MeCP2)^35,36^. We observed increased DAPI binding on cells expressing the *Sp*Swi6/*Sp*Clr4 proteins (Fig. 1b, c).

Second, and most striking, *Sp*Swi6/*Sp*Clr4 expression caused narrowing of the nucleus along the axis orthogonal to that defined by the nucleolus and Spindle Pole Body (SPB) (Fig. 1d). To observe this, we used a strain containing the nuclear pore complex (NPC) protein Nup49 and the nucleolus protein Nop1, labeled with GFP and mCherry, respectively. These results could be consistent with compaction of the chromatin in the Rabl configuration of chromosomes, with centromeres associated to the SPB clustered on one side of the nuclear envelope and telomeres located at the nuclear periphery at positions determined by the length of the chromosome arms (Fig. 1d)^38–43^. Consequently, compaction of the chromosomal arms between centromeres and telomeres induced by *Sp*Swi6/*Sp*Clr4 could result in the nuclear envelope being drawn inward, resulting in an ellipsoid nuclear shape with the long axis determined by the SPB-nucleolar positions^7,18,44^.

Finally, *Sp*Swi6/*Sp*Clr4 expression resulted in reduced stiffness, characteristic of increased chromatin compaction. Stiffening of the chromatin does work that drives separation of sister chromatids and partitioning of chromatin to different regions of the nucleus^45–47^. For example, it has been shown that depletion of nucleosomes that accompanies replication and mitosis results in stiffening of the chromatin as measured by an increase in effective spring constant (*k_s_*)^47^. The increase in *k_s_* is the consequence of the inverse relationship between *k_s_* and the local persistence length (*L_p_*) of chromatin, which increases with increasing chromatin density.

The mechanical stiffness and radius of confinement (*R_c_*) of chromatin can be measured by spatiotemporal tracking of individual chromatin regions with site-specific fluorescent probes integrated into the genome. Well-established relationships have been derived from polymer theory to calculate *k_s_* and *R_c_*from plots of mean-squared displacements (MSD) (See Methods, Equations 1 and 2)^45–47^. We chose to probe the spatiotemporal dynamics of the metabolic stress-response gene *INO1* locus. We chose this locus because we could measure two effects of *Sp*Swi6/*Sp*Clr4 expression: changes in chromatin stiffness and the adaptive response of *INO1* activation. It is known that nucleoplasm to nuclear envelope (NE) translocation concomitant with locus stiffening, results in more efficient *INO1* gene expression^48^.

To probe the locus spatiotemporal dynamics, we used a strain in which an array of 128 LacO sites is inserted into the genome adjacent to the *INO1* locus and the LacO-binding LacI protein fused to GFP is expressed. A bright fluorescent focus arising from the array-bound LacI-GFP can then be tracked. In this strain, NPC protein Nup49 was also labeled with GFP and Nop1 with mCherry as reference markers for the nuclear envelope and nucleolus, respectively (Fig. 2a, b). The MSD values for the *INO1*-LacI-GFP signal as a function of time were significantly higher for the activated state of the gene in the strain expressing *Sp*Swi6/*Sp*Clr4 compared to the reference strain also growing under activating conditions, and similar to repressed conditions for both types of cells (Fig. 2c). This indicated that in the presence of these heterochromatin forming proteins the locus behaved as if the cells were growing under repressed conditions. The values obtained for *R_c_* and *k_s_* also confirmed these results, with the gene showing a higher area of distribution (*R_c_*453,3 vs 315,4 nm) and lower effective spring constant (*k_s_* 0,2 vs 0,37 fN/nm) for cells expressing *Sp*Swi6/*Sp*Clr4 compared to reference cells, which means the locus is more flexible than expected in the activated state (Fig. 2d-g).

**Fig. 2.**
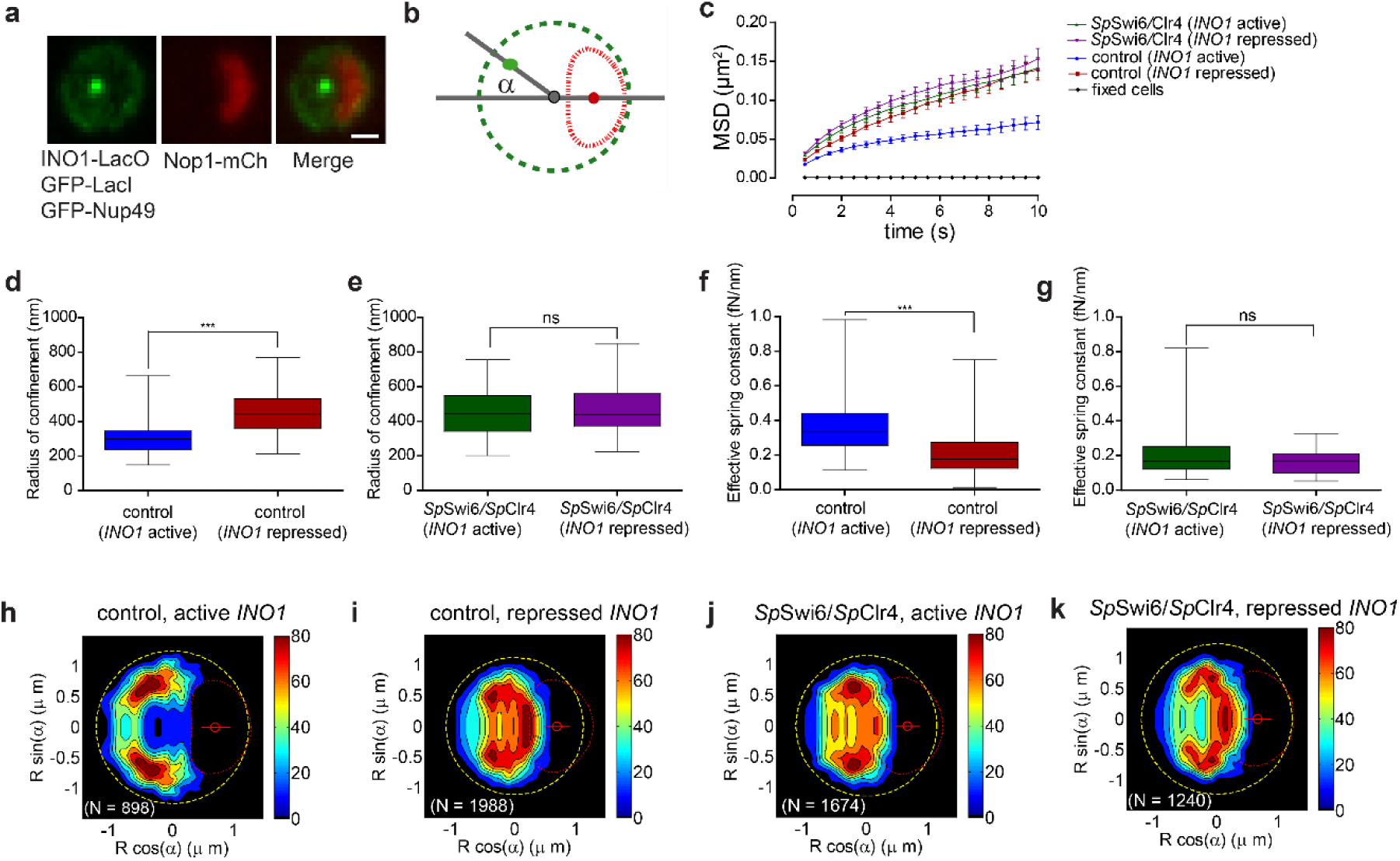
Active INO1 locus behave like condensed repressed chromatin in cells expressing SpSwi6/SpClr4 proteins. **a**, Representative images of a yeast strain nuclei in which an array of 128 LacO sites are integrated next to the INO1 locus and the LacO-binding protein LacI-GFP is expressed. Additionally, the NE and the nucleolus in these nuclei are labeled with GFP (Nup49) and mCherry (Nop1), respectively. Images shown are maximum projections of 250 nm Z stacks. Scale bar equals 1 µm. **b**, Positioning of the nuclear landmarks used to track the INO1 gene and to generate the statistical distribution maps for the localization of the INO1 locus. Green dashed circle (NE), red dashed ellipsoid (nucleolus), grey circle (nucleus center) and red circle (center of the nucleolus). The green circle represents the position of the gene in the nucleoplasm and the grey lines, the axes that connect the center of the nucleus with the center of the nucleolus and the center of the nucleus with the gene; α is the angle between these axes. **c**, MSD curves for the INO1 locus in reference and SpSwi6/SpClr4 strain under activating or repressed conditions, and for the gene in a population of fixed cells (black curve). **d**, Radius of confinement (R_c_) for the INO1 gene in an active state (blue) is significantly smaller than in the repressed state (red) for reference cells (two-sided t-test, p < 0.0001). **e**, R_c_ in the SpSwi6/SpClr4 strain under activating conditions (purple) does not show significant differences compared to the repressed state (grey). **f**, **g**, Spring constant (k_s_) for the INO1 activated gene in the reference (blue) significantly decreases compared to repressed cells (two-sided t-test, p = 0.0002), while there are no significant differences in R_c_ and k_s_ in the SpSwi6/SpClr4 strain under both conditions. **h**-**k**, Statistical maps for the INO1 localization obtained with nucloc software by superimposing nuclei of reference and SpSwi6/SpClr4-expressing cells grown under activating or repressed conditions. For these experiments, in addition to inositol conditions, cells were grown in SC-met. In the box and whiskers plots (d-g), the median is indicated as middle line, 25th and 75th percentile as boxes and the whiskers represent minimum and maximum values.

We have demonstrated that partitioning of the *INO1* locus to the NE from the nucleoplasm is accompanied by an increase in locus chromatin stiffness causing it to phase separate from the denser nucleoplasm chromatin to less dense perinuclear chromatin^48^. We therefore predicted that expression of *Sp*Swi6/*Sp*Clr4 could prevent this partitioning. We calculated maps of the statistical distribution of the *INO1* locus under repressed and activated conditions^43^. Under activating conditions (inositol starvation), the locus appeared to be confined to a region adjacent to the NE, but under repressed conditions (presence of inositol) the locus was broadly distributed throughout the nucleoplasm (Fig. 2h, i). We observed, however, a reduction of NE localization of the *INO1* locus in the strain expressing *Sp*Swi6/*Sp*Clr4 under activating conditions similar to the repressed state of the gene (Fig. 2j, k).

### *Sp*Swi6/*Sp*Clr4 expression increases genome stability to mutations

Genetic mutation is among the primary sources of genetic variation that determines rates of evolution, speciation and on short time scales, genome stability^49,50^. We next asked whether *Sp*Swi6/*Sp*Clr4 expression and resulting compaction of chromatin would cause a decrease in mutation frequency. We measured single nucleotide (SNPs) and insertion/deletion (indel) mutations with the widely used *CAN1* mutator assay, which is based on counterselection of the *CAN1* gene that encodes for the arginine permease Can1, an amino acid transporter responsible for the uptake of arginine from the environment^51–54^. Can1 also specifically transports the toxic arginine analog, canavanine. Consequently, we can select for individual clones containing *CAN1* mutations by growth on solid medium containing canavanine but not arginine (Fig. 3a)^52–54^.

**Fig. 3.**
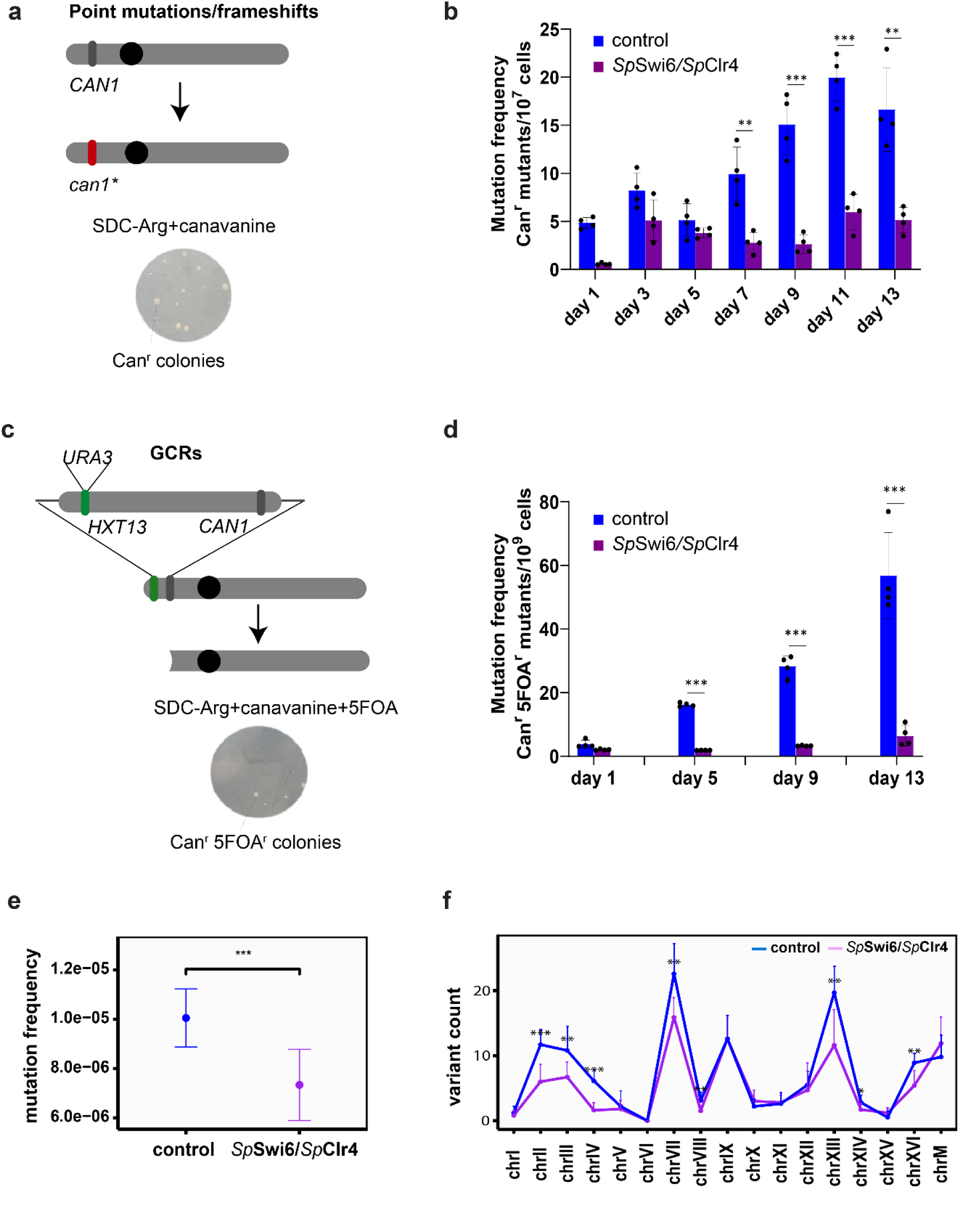
Genomic stability increases in SpSwi6/SpClr4 mutants. **a**, CAN1 mutator assay. Mutations in the CAN1 gene confer resistance to canavanine, a toxic arginine analog. Mutation frequency is quantified by measuring the frequency of canavanine resistant mutants on plates of synthetic medium lacking arginine and containing canavanine. **b**, Mutation frequency for the CAN1 gene in reference (DBY746) versus the DBY746-SpSwi6-SpClr4 strain, measured as Can^r^ mutants/10^7^ cells, on cultures incubated over 13 days. **c**, Gross Chromosomal Rearrangement (GCR) assay. The frequency of large chromosomal rearrangements is quantified by determining the frequency of mutations for two linked loci, URA3 and CAN1. Mutants are counter-selected on plates containing canavanine and 5-fluoroorotic acid (5-FOA), which is converted by the URA3 gene product to toxic 5-fluorouracil. **d**, GCR frequency, measured as Can^r^ 5-FOA^r^ mutants/10^9^, in reference (DBY746-URA3) versus the SpSwi6/SpClr4 strain, for cells cultured over 13 days. Experiments were performed in four biological replicates. **e**, mutation frequency in reference (DBY746) versus the DBY746-SpSwi6-SpClr4 strain after 13 days incubation. **f**, variant count per chromosome in reference (DBY746) versus the DBY746-SpSwi6-SpClr4 strain after 13 days incubation. For these assays cells were cultured in SC-met before being plated in the corresponding selection plates mentioned above. Error bars represent standard deviations and asterisks represent the significance of the P-values calculated with Student’s t-tests.

To characterize the type and quantify the frequency of mutations that occur in reference versus *Sp*Swi6/*Sp*Clr4-expressing cells, we monitored the frequency of Can^r^ mutations (number of colonies) over several days of incubation (Fig. 3b). We used as reference the *S. cerevisiae* DBY746 strain that has been used before for these types of studies, and for which it has been observed that the number of mutants with Can^r^ increases rapidly, after several days of growth^55^ (Methods section: ‘Mutation frequency measurements’). For the first 5 days of incubation, we did not observe a significant difference in the frequency of mutants for reference or *Sp*Swi6/*Sp*Clr4-expressing strains although the values were always higher for the reference strain. However, after day 7 the difference became significant, and the resulting frequencies of mutation were 3-4 times lower for the *Sp*Swi6/*Sp*Clr4-expressing than reference cells by day 13 (Fig. 3b).

We also analyzed the type of mutations that appeared to confer resistance to canavanine for 10 of the Can^r^ mutants (Extended Data Table 1). For one of the clones obtained from the reference strain, we were not able to amplify the *CAN1* locus which suggests that a large chromosomal rearrangement might have occurred in this clone. For the rest of the clones from reference, sequencing of the *CAN1* gene showed that six contained single base substitutions, three single deletions and one exhibited a triple deletion (Extended Data Table 1). Three of the clones contained two or more types of mutations. For the base substitutions four were transversions C → A or G → C, and four were transitions G → A or A → G. The deletions were ΔC or ΔT. For the DBY746 *Sp*Swi6/*Sp*Clr4-expressing strain, we were able to amplify and sequence the 10 colonies selected, indicating no chromosomal rearrangement. Additionally, all colonies exhibited only one single point mutation per strain. Most of the mutations observed were base substitutions present in eight of the 10 clones. The other two strains contained a single base deletion ΔC in both cases. The base substitutions were mainly transversions the type C → A, G → C, G → T and a single transition C → T (Extended Data Table 1).

We used a second approach to determine the frequency of gross chromosomal rearrangement (GCR) based on an assay in which two counter-selectable markers are linked and the loss of both is measured by growing the cells in the presence of a double selection medium. In this case the markers are, *CAN1* and *URA3*, with *URA3* placed at the *HXT13* locus, which position it nearby the *CAN1* gene. The strains to analyze are grown in medium containing canavanine and 5-fluoroorotic acid (5-FOA)^56,57^. The uracil biosynthesis pathway can convert the nontoxic 5-FOA into toxic 5-fluoro-uracil and the *URA3* gene encodes for a product that catalyzes a key step in this synthesis. The treatment of cells with both canavanine and 5-FOA will thus select for *CAN1-* and *URA3-* inactive mutants. To avoid loss of cell viability by rearrangement or deletion of the whole cassette flanked by *CAN1–URA3*, these markers are usually inserted into a position proximal to the telomere in the chromosome arm (Fig. 3c). The chances that point mutations can inactivate two different markers simultaneously is below the level of detectability. Consequently, unless cells are also treated with DNA mutagens, the *CAN1–URA3* reporter likely measures GCRs^56–59^. The types of rearrangements that are expected to be detected with this assay include different inter- and intra-chromosomal fusion events as well as broken chromosomes repaired by *de novo* telomere additions^56–60^.

We measured the GCR frequency for the reference DBY746 and DBY746-*Sp*Swi6-*Sp*Clr4 grown for a period of 13 days. After day 5 there was a significant increase in GCR frequency for the reference compared to the *Sp*Swi6/*Sp*Clr4-expressing strain that reached 10-fold by day 13 (Fig. 3d). Overall, it appears that the expression of *Sp*Swi6/*Sp*Clr4 and consequent compaction of chromatin results in a significantly decreased number of mutations in the genome, including large chromosomal rearrangements.

Finally, we analyzed mutation frequency for the whole genome in the DBY746 strain expressing *Sp*Swi6 and *Sp*Clr4 compared to reference and in the absence of any selection pressure. For this, we performed whole genome sequencing (WGS) on 10 isolated single colonies for the reference or the *Sp*Swi6/*Sp*Clr4 expressing cells after 13 days incubation.

Similar to the estimation by *CAN1* selection, the mutation frequency in the reference strain was higher than that of the *Sp*Swi6/*Sp*Clr4-expressing strain (approximately 7.3 versus 10 per one million bases) (Fig. 3e). Most mutations, in both strains, occurred on chromosomes II, III, IV, VII, VIII, XIII, and XVI (Fig. 3f). The variants were primarily indels (approximately 73 for reference and 100 for SpSwi6/SpClr4 strains) rather than point mutations (approximately 16 for reference and 20 for SpSwi6/SpClr4 strains). Indel mutations were found mostly on chromosomes II, IV, and XIII, as well as more point mutations in chromosomes III and XVI compared to *Sp*Swi6/SpClr4-expressing cells (Extended Data Fig. 1).

### Global meiotic recombination rates decrease in *Sp*Swi6/*Sp*Clr4 expressing cells

Meiotic recombination is another important driver of genetic variation, and a key source of genome evolution in organisms that reproduce sexually. This process provides new allelic combinations that serve as pools for natural selection as well as artificial selection of high-performing genotypes for industrial purposes^61^. *S. cerevisiae* is unique in that its rate of recombination is so high that it can reach nearly 100 % linkage equilibrium within 6 generations of inbred crosses^62–66^. Indeed, mitotic recombination between artificially constructed ectopic repeats can occur as efficiently as allelic recombination in *S. cerevisiae*^62^. However, for example, in *S. pombe*, ectopic recombination has been reported to take place much less frequently (10 – 1000 times) than allelic recombination^67,68^. Among eukaryotic organisms, recombination rates vary inversely with genome size over several orders of magnitude but for *S. cerevisiae* it is 3 times higher than predicted^65,69–71^. We next asked whether the expression of *Sp*Swi6/*Sp*Clr4 would affect rates of meiotic recombination.

To quantify the recombination rate for *S. cerevisiae* we used a high-throughput and low-cost strategy in which three fluorescent marker protein-coding genes are integrated into chromosomes in different positions and flow cytometry is used to measure recombination rates and crossover patterns, based on compositions of fluorescent proteins expressed in spores^61^. The fluorescent protein-coding genes were integrated at optimal distances from each other in a particular chromosome to measure recombination rates^61^. These strains were crossed with non-marker strains to obtain diploids that would produce spores with different rearrangement patterns depending on the recombination events that occur in the chromosomes (Fig. 4a). By calculating the frequencies of these events per segment, we were able to quantify the values for rate of recombination between each different pair of markers in each strain (See Methods, Equation 3).

**Fig. 4.**
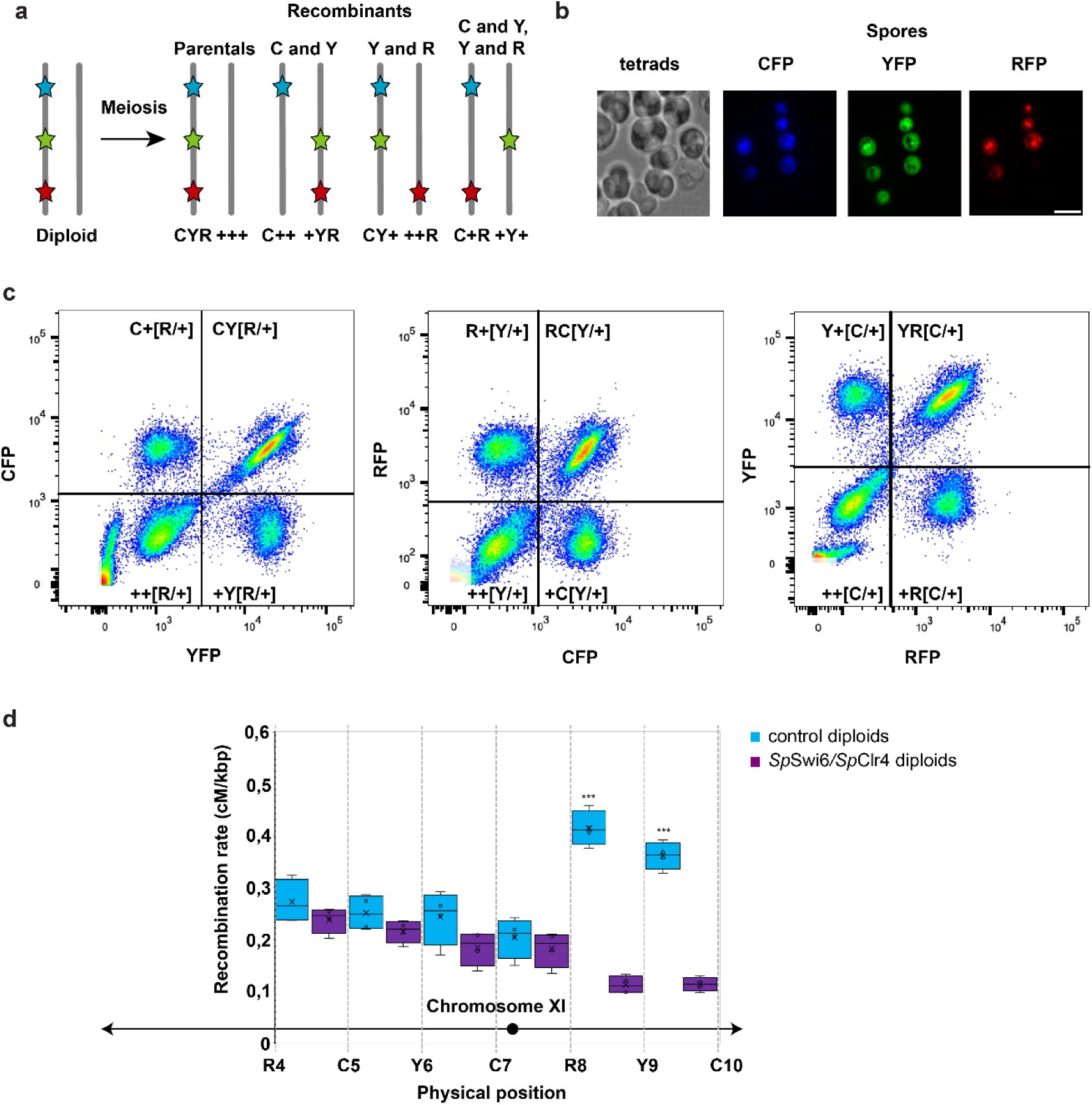
Global recombination rate decreases in S. cerevisiae cells expressing SpSwi6/SpClr4 heterochromatin. **a**, Meiotic chromosome segregation for three linked markers. Spores were classified into different classes according to the pattern of the markers they contained by using the three letters C (cerulean), Y (yellow), and R (red). The symbol ‘+’ was used to indicate the absence of a particular marker. Thus, for a distribution pattern CYR, parental spores were denominated CYR and +++, recombinant spores between C and Y, or between Y and R, were designated as C++ and +YR, or CY+ and ++R, respectively. Finally, double recombinant spores were named C+R and +Y+. **b**, Imaging of tetrads and fluorescent spores following sporulation and spore isolation, respectively. Scale bar represents 5 µm. **c**, Examples of projections obtained for one sample representing the fluorescent intensities for the different pairs of markers: Venus (YFP) versus Cerulean (CFP), Cerulean (CFP) versus mCherry (RFP), and mCherry (RFP) versus Venus (YFP), respectively. Indicated in each quadrant per projection are the classifications of the spores, with the symbol [X/+] representing that the designated marker may be present or not at this position. **d**, Recombination rates for the different tri-fluorescent tester and SpSwi6/SpClr4 containing diploids along chromosome XI of S. cerevisiae. The centromere of the chromosome is indicated with a black circle and the extremes with black arrows. Four biological replicates were used to calculate these results. Asterisks represent levels of significance for the P-values obtained from Student’s t-tests between strains. Median is indicated as middle line, average as a black X, 25th and 75th percentile as boxes and whiskers represent minimum and maximum values.

We transformed the strains containing the three fluorescent markers spanning a total of 7 different consecutive sites in chromosome XI with the *Sp*Swi6/*Sp*Clr4 expression cassette. Thus, we used three strains, each expressing three markers in three consecutive sites that span the region containing the 7 sites. For example, strain 1 expressed markers in sites R4C5Y6, strain 2 Y6C7R8, and strain 3 R8Y9C10; with R (RFP), C (CFP), Y (YFP), being code for the color of the fluorescent marker expressed. These tri-fluorescent strains expressing *Sp*Swi6/*Sp*Clr4 and their parents were crossed with either *Sp*Swi6/*Sp*Clr4-integrated or reference W303 strains, respectively, to obtain diploids containing either two or none *Sp*Swi6/*Sp*Clr4 cassettes. These diploids were induced to sporulate, followed by spore isolation, and finally FACS analysis of the spores (Fig. 4b, c). Each spore was classified according to its fluorescence pattern, which allowed to calculate values of recombination rates for all consecutive sites in each strain (Fig. 4d, See Methods, Equation 3). Recombination rates ranged from 0.26 to 0.41 cM/kbp among the 6 regions on chromosome XI for the reference diploids, and from 0.11 to 0.23 cM/kbp for the *Sp*Swi6/*Sp*Clr4 containing diploids (Fig. 4d). In both cases higher values of recombination rate were observed towards the end of the chromosome arm as expected^72^. On average, the values obtained for reference strains are within the range previously reported for *S. cerevisiae* ^61,65^, however the averages for strains expressing the *Sp*Swi6/*Sp*Clr4 factors are almost two times lower. This difference is mainly due to large decreases in the rates of recombination in the peripheral right arm of the chromosome, but all regions showed decreases (Fig. 4d).

### *Sp*Swi6 pervasively binds to the genome

To explore the topology of *Sp*Swi6 and *Sp*Clr4 interactions with the chromosomes of *S. cerevisiae*, we performed ChIP-seq of Flag-tagged *Sp*Swi6 in *Sp*Swi6-5Flag/*Sp*Clr4-expressing cells. A similar strain where *Sp*Swi6 does not harbor the Flag tag was used to control for the specificity of the anti-Flag antibody and a strain expressing *Sp*Swi6-5Flag but no *Sp*Clr4 was included to assess the contribution of H3K9 methylation on *Sp*Swi6 occupancy. A small amount of chromatin from 3Flag-Swi6 *S. pombe* cells was spiked in the chromatin prior to the immunoprecipitation for normalization. We found that *Sp*Swi6-5Flag binds pervasively across the whole genome (Fig. 5 and Extended Data Fig. 2a), unlike the more localized distribution of binding found on *S. pombe* chromosomes mainly at pericentromeric and sub telomeric DNA regions, and the mating type (mat) locus^73^. The pervasive binding Swi6 in *S. cerevisiae* most likely is the consequence of the lack of targeting activities in this organism. In *S. pombe*, the Swi6/Clr4 system is targeted/nucleated by at least two mechanisms: an RNAi-based mechanism and a mechanism involving the MTREC complex. These components are absent in *S. cerevisiae* so Swi6 is left unguided^74^. The no-tag control strain only shows signal on the open reading frame of highly expressed genes as often observed in ChIP experiments^75–77^. Surprisingly, *Sp*Swi6-5Flag occupancy was virtually identical in cells not expressing *Sp*Clr4. ChIP may not be sensitive enough to pick up more subtle but functionally essential effects on binding, such as decreased exchange rates of *Sp*Swi6 with chromatin containing H3K9me. This argument is further supported by the less even distribution of mutations among the chromosomes, implying different degrees of compaction. Paradoxically, *Sp*Swi6-5Flag binding is highest in nucleosome-free regions found in promoters (Extended Data Fig. 2b). This again, may suggest that the interactions of *Sp*Swi6 with H3K9me are more subtle, but functionally essential. To assure, however, that *Sp*Clr4 is active when expressed in *S. cerevisiae*, we directly measured H3K9 methylation. Both H3K9 di- and tri-methylation (H3K9me2/3) were detected in association with *Sp*Swi6 in co-immunoprecipitation experiments from *Sp*Swi6-5Flag/*Sp*Clr4-expressing *S. cerevisiae* cells (Extended Data Fig. 2c), confirming that the *Sp*Clr4 methyltransferase is active and that *Sp*Swi6 can bind to methylated H3K9. The methylation was low, but detectable in the Inputs. However, ChIP-seq experiments for H3K9me2 in *Sp*Swi6-5Flag/*Sp*Clr4 and *Sp*Swi6-5Flag-expressing cells did not result in detectable signal over background (Extended Data Fig. 2a). Because we only measured H3K9me2, we cannot rule out the possibility that either H3K9me1 or H3K9me3 accumulates on chromatin in *Sp*Clr4-expressing cells and accounts for the phenotypic effects of *Sp*Swi6/*Sp*Clr4 expression. In fact, lately it has become more evident that H3K9me2 and H3K9me3 can establish two different chromatin states, with H3K9me3 domains linked to high levels of HP1 binding and low levels of transcription, while H3K9me2 appears to be related to lower levels of HP1 recruitment and a more transcriptionally permissive state^78,79^.

**Fig. 5.**
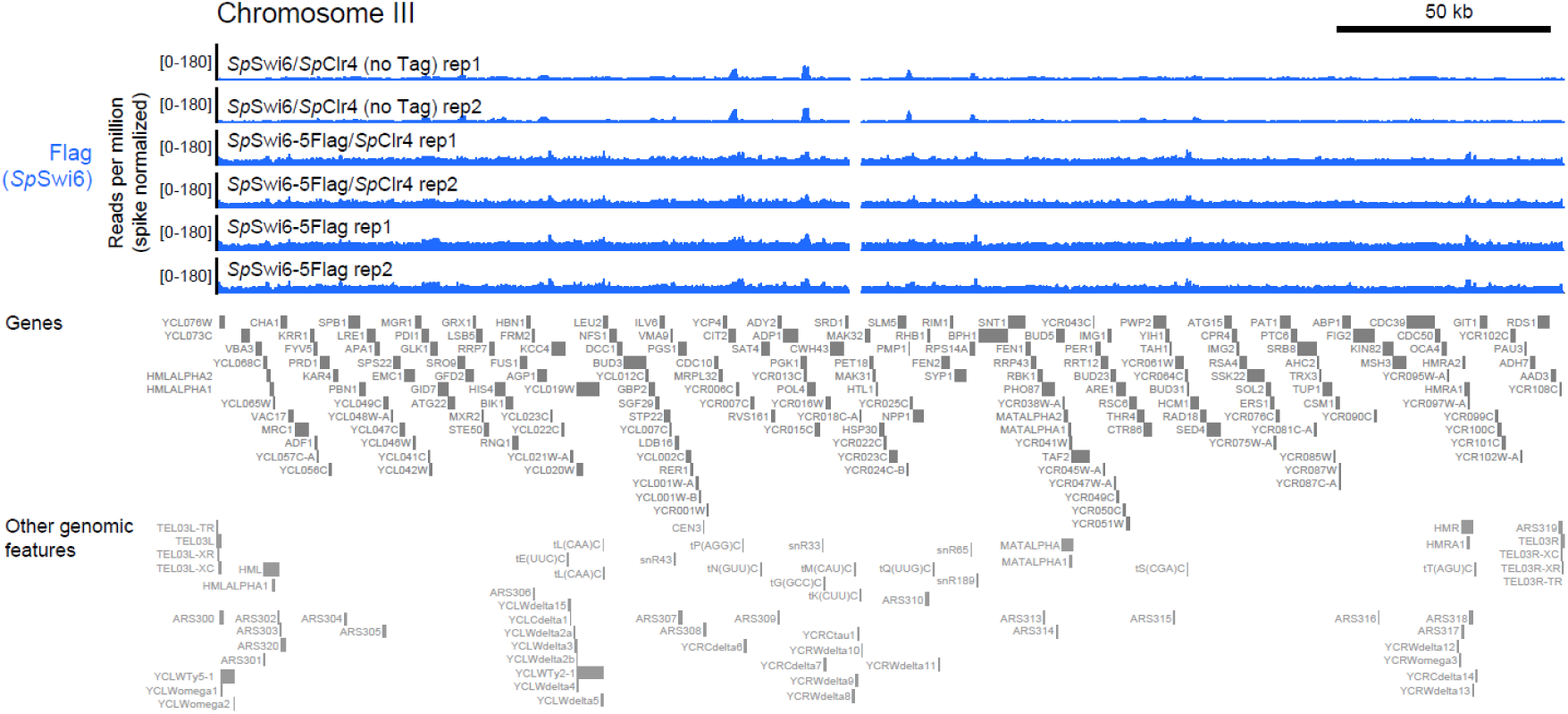
SpSwi6 binds pervasively across the genome when expressed in S. cerevisiae. A genome browser view of anti-Flag (SpSwi6) ChIP-seq over the S. cerevisiae chromosome III, representative for the whole genome data set. S. pombe-spike normalized read counts are shown for both replicates from ChIPs performed in S. cerevisiae cells expressing SpSwi6/SpClr4 (no tag control), SpSwi6-5Flag/SpClr4 or SpSwi6-5Flag (no SpClr4 control). Genes and other genomic features are shown below the data tracks.

### *Sp*Swi6/*Sp*Clr4 expression stabilizes a metabolically engineered strain

We next asked whether the genome compaction and stabilization to mutation caused by *Sp*Swi6/*Sp*Clr4 expression could help stabilize metabolically engineered strains. There is a great need to improve the genetic stability of *S. cerevisiae* strains engineered for chemical production and other biotechnological applications. Strains engineered with heterologous metabolic pathways that happen to cause growth defects or compete with cell division are particularly prone to mutations that inactivate or diminish their productivity. Dynamic controls can help improve production and preserve strain stability by dividing cell cultures into a growth phase dedicated to allowing essential pathways to drive cell division, and a production phase in which resources are rerouted towards chemical production^80^. However, these controls can also suffer from mutations that override the engineered growth regulatory systems. For example, ethanol is a major unwanted by-product of *S. cerevisiae* fermentations designed to produce other chemicals. Therefore, limiting ethanol production improves desired product yields. However, ethanol fermentation is also required for growth and division of yeast on glucose and thus splitting this process into a dynamic two-phase fermentation, one dedicated for cell growth and another for chemical production, has been shown to be advantageous (Fig. 6a)^81^. Thus, to address the competition between growth (and its associated ethanol by-product formation) and production of pyruvate-derived chemicals, we previously developed dynamic controls using optogenetics to regulate the transcription of *PDC1* pyruvate decarboxylase^81^ (Fig. 6b). These controls enable cell growth in blue light and, when sufficient biomass is accumulated, redirection of pyruvate towards other products of interest by switching to darkness, thereby reducing cell growth and ethanol formation^81^. However, long incubations (40-100 hours) of such strains in non-permissive, dark, conditions can lead to “cheating” cells that escape optogenetic regulation.

**Fig. 6.**
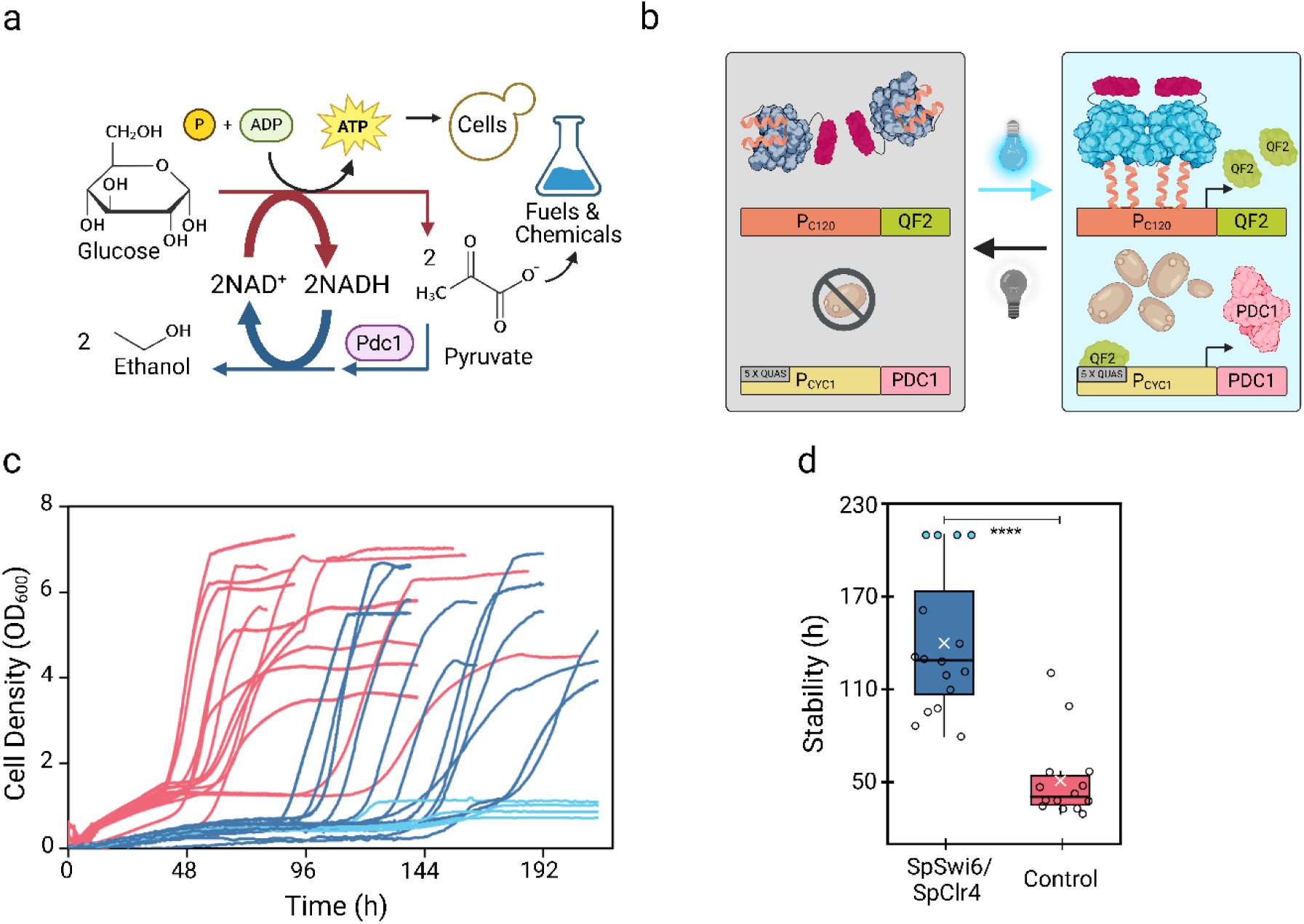
SpSwi6 and SpClr4 extend genetic stability of light dependent growth in S. cerevisiae. **a,** Pdc1p is essential for S. cerevisiae growth on glucose and ethanol production, as it is required to generate acetaldehyde, the electron acceptor needed to regenerate the NAD^+^ reduced during glycolysis. This results in ethanol by-product formation and consumes most of the pyruvate needed to produce other fuels and chemical of interest. **b,** Circuit design of OptoQAmp2, used to control the expression of PDC1 in S. cerevisiae^82^. OptoQAmp2 utilizes the VP16-EL222 transcription factor from E. litoralis^84^, dimerizes in response to blue light to activate its cognate promoter P_C120_ and express QF2, a strong transcriptional activator from Neurospora crassa, which in turn activates expression of PDC1 from its own cognate promoter, 5xQUAS-P_CYC1_. As a result, this strain depends on blue light to express PDC1 and is thus unable to grow on glucose in darkness. **c,** Replica growth curves of this optogenetic strain expressing SpSwi6 and SpClr4 (blue, n=16) or carrying an empty vector as a control (red, n=14) cultured in glucose and dark (non-permissive conditions). Expressing SpSwi6 and SpClr4 increases the strain stability 2.6-fold compared to the control; in fact 4 replicas remained stable for the duration of the experiment (cyan, 214 h). **d,** Stability distribution of optogenetic strain expressing SpSwi6 and SpClr4 (n= 16, blue and cyan) shows a 2.6-fold increase in stability (**** p<8.78 × 10^-7^) relative to the control strain (n = 14, red). Box plot legend: circles are individual data points, white X represents the average, horizontal line the median, cyan-filled circles in (d) correspond to the 4 replicas that remained stable throughout the experiment, asterisk above boxes indicate statistical significance determined by a two-sided t-test (α=0.05, p<8.78 × 10^-7^, f see methods).

To explore the effect of *Sp*Swi6/*Sp*Clr4 expression on the stability of engineered yeast, we expressed them in an optogenetically controlled strain that invariably loses optogenetic control or “cheats” under non-permissive dark conditions. We integrated the cassette expressing *Sp*Swi6/*Sp*Clr4, controlled by the repressible TetO and *MET17* promoters respectively, into a strain containing the blue light-responsive OptoQAmp2 optogenetic circuit^82^ to control expression of *PDC1* in a triple *PDC* background (*Δpdc1, Δpdc5, Δpdc6*), (Extended Data Tables 2 and 3). We then compared the time and frequency of cheating of this strain when incubated in non-permissive conditions (darkness) against a control strain of identical genetic background but lacking *Sp*Swi6 and *Sp*Clr4 (Extended Data Table 2). In both strains, we also introduced a blue fluorescent protein reporter (mTAGBFP2) gene to distinguish cheating cells from possible contaminants.

We conducted these experiments using ChiBio minibioreactors, which automatically monitor optical density in real time and can illuminate cultures with blue light (457 nm) on any light schedule throughout cultivation^83^. Cultures were inoculated at a starting cell density of OD_600_=0.01 and incubated for several days. In permissive conditions (blue light), both strains (in 3-4 replicates) grow with similar lag times, although the strain expressing *Sp*Swi6/*Sp*Clr4 showed a slower growth rate (0.13 h^-1^ vs 0.23 h^-1^) and lower maximum biomass accumulation than the control strain (Extended Data Fig. 3a). This growth defect, however, seems to be specific to the optogenetically controlled strain, as no defect was observed when expressing *Sp*Swi6/*Sp*Clr4 in the reference CEN.PK2.1C strain (Extended Data Fig. 3b). To test whether *Sp*Swi6/*Sp*Clr4 expression improves the stability of engineered strains, we incubated cultures inoculated to a cell density of OD_600_=0.01 in non-permissive conditions (darkness) for 214 hours and measured the frequency and time at which cheaters started to grow in the dark. All 14 replica cultures (100%) of the control strain escaped the optogenetic growth regulation at random times within ∼120 hours of cultivation, as early as ∼37h from the start of cultivation, and with an average stability of 55 ± 29 h (Fig. 6c). In contrast, of the 16 replica cultures of the strain expressing *Sp*Swi6/*Sp*Clr4, the first cheater did not appear until almost 52h (Fig. 6c red lines) and 144h (Fig 6c dark blue lines) after starting the dark incubation, and cultures remained stable on average of 144 ± 24 h, a 162% increase in average strain stability (Fig. 6c and 6d). Moreover, 4 replica cultures of this strain (25%) remained stable for the entire duration of the experiment (214 h). When exposing these 4 remaining cultures to blue light, they all started to grow, (Extended Data Fig. 4), confirming the strains had remained alive and stably regulated. This 2.6-fold improvement in average strain stability brought by *Sp*Swi6/*Sp*Clr4 is highly statistically significant (*p* value < 8.78 × 10^-7^).

## Discussion

How can expression of *Sp*Swi6 and *Sp*Clr4 result in global changes of chromatin density and motion that impact genome-wide mutation and recombination rates? Results linking mutation rate to chromatin compaction varies depending on which cells are studied and the type of mutation^85,86^. For instance, in mouse embryonic fibroblasts, knockouts of all six SET domain lysine methyltransferase genes lacked all H3K9 methylation states, derepressed nearly all families of repeat elements and displayed genomic instabilities. Furthermore, mutant cells no longer maintained heterochromatin organization and lost all electron-dense heterochromatin^87^. The relationship between recombination rate and chromatin compaction is clearer, where open euchromatin exhibits higher recombination rates than compacted heterochromatin^88^. These observations do not, however, provide mechanistic insight into the relationship between chromatin dynamics and genome stability.

It has been shown that the fidelity of biological processes can be achieved through stochastic molecular dynamics in several cellular processes. For instance, Nicklas proposed that the stochasticity of microtubule dynamics were the critical feature in achieving high fidelity chromosome segregation^89^. The dynamics that we observe in chromatin is due to ATP-dependent processes, such as cohesin-driven loop extrusion^90–93^. Consequently, chromatin loops are constantly fluctuating in volume (expanded to contracted), physical size, and density (number of loops per kilobase pair). The increased chromatin dynamics that we observe in *Sp*Swi6/*Sp*Clr4-expressing *S. cerevisiae* could more easily expose it to DNA repair machinery, thus suppressing mutations such as occurs with the assembly of RecA^94,95^. RecA assembles into protein filaments on a single-stranded DNA (ssDNA), which appears as an intermediate structure at a double-strand break sites, forming kilobase-long presynaptic filaments that mediate homology search and strand exchange reactions^96^. This feature of RecA assembly dynamics is conserved in Rad51, the RecA ortholog in yeast. The process of filament assembly establishes a kinetic proofreading cascade that enables the cell to mount an SOS DNA repair response to exposed ssDNA^94,95^. RecA proofreads the ssDNA through its binding fluctuations. There is a timescale of nucleation, dependent on RecA concentration, and a second time-scale dependent on the ssDNA fluctuation (i.e. the tendency of ssDNA to adopt a random coil). At low RecA concentration, nucleation events will be rare, and filaments will not assemble. If fluctuations are too rapid, ssDNA will collapse, and filaments will not assemble. At a low rate of chromatin extrusion, however, the chromatin may also not be exposed long enough for filaments to assemble. Tuning the timescales of nucleation and ssDNA fluctuation is therefore critical to filament assembly and SOS DNA response.

Why do we observe more indels than point mutations? In many organisms, point mutations are more common than indels. This trend is typically due to the higher likelihood of single base changes during DNA replication compared to the more complex processes required to insert or delete bases. However, the idea that point mutations are more frequent than indels across all contexts is not a general rule. Several factors that could affect elevated indels are replication slippage (repeat sequences (microsatellites) where the DNA polymerase can slip), difficulty in NHEJ DNA repair, mobile genetic elements, structural features of the DNA (hairpin quadruples), biased gene conversion (preference over one strand of the genome over another during replication), and environmental factors. We could speculate that during early (initial) stages of adaptation (which is comparable to our experiment here), major genomic rearrangements indels are a major source of adaptive mechanism, eventually being replaced by point mutations (as observed in many lab evolution experiments) ^97–99^.

Stress is known to increase mutation rates in microorganisms^100^. Therefore, the enormous selection pressure on our light-dependent strain to escape optogenetic regulation when incubated in darkness for extended periods was ideal to study the effects of *Sp*Swi6/*Sp*Clr4 on the genetic stability of engineered yeast. The random nature of the times at which isogenic cultures adapt to grow in the dark supports the hypothesis that these adaptations occur by mutation, which we confirmed by the complete loss of light regulation observed when re-growing the cheater cells (Extended Data Fig. 5 and Supplementary Note 1). Long incubations in the dark impose on this strain a higher selection pressure for destabilizing mutations than the typical heterologous biosynthetic pathway, as darkness for these strains is essentially toxic. Moreover, genetic instabilities can be easily detected and quantified in this model strain simply by measuring the lag phase of cultures incubated in the dark, which would be more laborious and imprecise by monitoring reduced productivity in strains engineered for chemical production. Therefore, the remarkable 162% increase in genetic stability achieved in this model strain demonstrates the great potential of *Sp*Swi6 and *Sp*Clr4 to improve the stability of biotechnologically relevant engineered strains.

Finally, the harsh conditions in which *S. cerevisiae* is maintained in industrial practices make engineered strains susceptible to genetic instability including chromosomal rearrangements, causing the loss of beneficial traits or the appearance of phenotypes such as cell aggregation, harmful for production^101–104^. In fact, *S. cerevisiae* isolates from different industrial sources show high genomic intra-strain variability^105,106^. Consequently, the increased, or effectively completely stable maintenance of an engineered strain that we achieve through expression of *Sp*Swi6/*Sp*Clr4 could be a game-changer, as genetic instability is a major limitation to the long-term productivity of biotechnologically relevant strains^104,107,108^.

## Acknowledgements

The authors thank Geeta J. Narlikar (University of California, San Francisco), Jason Brickner (Northwestern University), Matt Kaeberlein (University of Washington), Matthieu Falque (Université Paris-Saclay, France), and Danesh Moazed (Harvard Medical School) for providing reagents, plasmids, and strains.

## Funding

This work was supported by Canadian Institutes of Health Research (CIHR) grant MOP-GMX-152556 and Human Frontiers Science Program grant RGP0034/2017 (S.W.M). The section on ChIP-seq was in part funded by Canadian Institutes of Health Research (CIHR) grant PJT-156383 (F.R.). The section on stability of optogenetic strains (J.L.A) was funded by the DOE Center for Advanced Bioenergy and Bioproducts Innovation (U.S. Department of Energy, Office of Science, Office of Biological and Environmental Research under Award Number DE-SC0018420), as well as a separate award by the same agency and office (Award Number DE-SC0022155). Any opinions, findings, and conclusions or recommendations expressed in this publication are those of the authors and do not necessarily reflect the views of the U.S. Department of Energy.

## Author contributions

Conceptualization: L.G. and S.W.M.; Methodology: L.G., S.A.G.E., C.J., M.G, C.P.; Data Analysis-Writing-Review: L.G., S.A.G.E., C.J., C.P., M.G, A.S., K.B., F.R., J.L.A., S.W.M.

## Declaration of interests

SWM is a shareholder in Aarvik Therapeutics and member of the Scientific Advisory Board, EPOK Therapeutics.

## Methods

### Strains and plasmids

To analyze the effects of *Sp*Swi6 and *Sp*Clr4 expression on genome compaction of *S. cerevisiae,* two plasmids were generated (Extended Data Table 2). Briefly, plasmids containing *Spswi6* and *Spclr4* sequences (pET30a-*Sp*Swi6 and 2CT[1]-*Sp*Clr4) were provided by Geeta J. Narlikar at the University of California, San Francisco (Extended Data Table 3). *Spswi6* was subcloned into pCM189 under the control of the CYC1 promoter and a tet operator array containing 7 tetO sites, while *Spclr4* was subcloned into a p413-ADH1 vector where the ADH1 promoter had been previously replaced by a *MET17* promoter. Both pCM189-tetO7-*Sp*Swi6 and p413-MET-*Sp*Clr4 or empty plasmids were transformed in a W303 strain (Extended Data Table 3) and used to quantify DAPI signal and relative chromatin compaction.

To analyze nuclear morphology, a strain (TMS1-1A) containing a plasmid (pASZ11-NupNop) expressing GFP-Nup49 and mCherry-Nop1 to mark the NE and the nucleolus, respectively, was imaged as control^109^. This strain was also transformed with the plasmids previously described (Extended Data Table 2).

To generate an integration cassette containing both tetO7-*Sp*Swi6 and MET-*Sp*Clr4 expression cassettes, that could be inserted into different strains, the backbone of the yeast integration plasmid YIp204-PADH1-atTIR1-9myc was used (Addgene, Plasmid #99532) (Extended Data Table 3). In this case, any cassette subcloned into this plasmid can be inserted into the genome of *S. cerevisiae* by linearizing the plasmid at the *TRP1* locus (Bsu36I site) and then transforming the linearized vector into a tryptophan auxotroph strain. Briefly, the sequence that encodes for the TIR1-9myc protein was first removed by reverse PCR and then both tetO7-*Sp*Swi6 and *MET17-Sp*Clr4 expression cassettes were cloned consecutively at this site by using the GeneArt Seamless Cloning and Assembly kit (ThermoFisher Scientific) (Extended Data Table 3). By employing the same strategy, a 5Flag tag was inserted at the coding sequence for the C-terminus of *Sp*Swi6 to allow for detecting the expression of the protein by Western blot, co-immunoprecipitation and ChIP experiments (Extended Data Fig. 6 and 7). The plasmid generated (YIp204-tetO7*-Sp*Swi6-5Flag-*MET17*-*Sp*Clr4) was then linearized at the *TRP1* locus, purified, and stored for transformation (Extended Data Table 3).

To quantify the statistical distribution and mechanical properties of the *INO1* gene, the LMY52 strain^110^ (Extended Data Table 2) was employed as control (provided by Jason Brickner, Northwestern University, USA). This strain contains an array of 128 LacO sites fused to the *INO1* locus and expresses LacI-GFP and NE and nucleolus proteins fused to fluorescent proteins (GFP-Nup49 and mCherry-Nop1). This strain was transformed with the YIp204-tetO7-*Sp*Swi6-5Flag-*MET17*-*Sp*Clr4 or a non *Sp*Swi6/S*p*Clr4 linearized plasmid.

The *S. cerevisiae* DBY746 strain (Extended Data Table 2) (provided by Matt Kaeberlein, University of Washington, USA) was used as background to measure mutation frequency driven by small base substitutions/deletions/insertions. To assay the effect of *Sp*Swi6/*Sp*Clr4 expression on mutation frequency, the integration plasmid YIp204-*tetO7-Sp*Swi6-5Flag-MET*-Sp*Clr4 was transformed into this strain. To measure large chromosomal rearrangements in the DBY746 and DBY746-*Sp*Swi6-S*p*Clr4 strains, both were transformed with a *URA3* cassette designed with flanking homologous regions to replace the *HXT13* locus situated 7.5 kb upstream of the *CAN1* locus.

For recombination rate measurements, tri-fluorescent tester strains (Extended Data Table 2): 348 (SK1-XI-R4C5Y6), 400 (SK1-XI-Y6C7R8), 369 (SK1-XI-R8Y9C10) were provided by Matthieu Falque at Université Paris-Saclay, France^61^. These strains containing different arrangements of three-colored fluorescent protein-coding genes (Cerulean, Venus and mCherry) in chromosome XI were crossed with a W303-*MAT*a strain previously transformed with the empty YIp204 integration plasmid (Extended Data Table 2) to generate control diploids to use for sporulation in the recombination rate assays. The tri-fluorescent testers and W303-*MAT*a haploids were transformed separately with the plasmid YIp204-tetO7*-Sp*Swi6-5Flag-*MET17*-*Sp*Clr4 to integrate *Spswi6/Spclr4* into their genomes. Next, each tri-fluorescent tester strain expressing *Sp*Swi6/*Sp*Clr4 proteins was crossed with the W303-*MAT*a strain also expressing these proteins to obtain *Sp*Swi6/*Sp*Cl4 diploids for the recombination tests (Extended Data Table 2).

ChIP-seq was performed in the W303-*Sp*Swi6-5Flag/*Sp*Clr4 strain previously generated. As a control, a similar strain with no Flag tag was generated by integrating plasmid YIp204-tetO7*-Sp*Swi6-*MET17*-*Sp*Clr4. Also, a strain expressing *Sp*Swi6 but no *Sp*Clr4 was generated by integrating plasmid YIp204-tetO7*-Sp*Swi6-5Flag. Finally, an *S. pombe* strain expressing a 3Flag-Swi6 protein was used as a spike-in control in all ChIP samples (Extended Data Table 2).

Plasmids for optogenetic strains were constructed with restriction-based cloning and Gibson assembly with enzymes acquired from New England Biolabs (Ipswich, MA, USA). *S. cerevisiae* strains generated in this study were transformed by electroporation, as previously described^111^. Unless specified, all strains were grown and maintained in SC supplemented with 3% v/v glycerol and 2% v/v ethanol (SC-GE), and the corresponding dropout media for selection of transformants with auxotrophic markers. We used a triple *PDC* knockout strain (*Dpdc1, Dpdc5, Dpdc6,* called SGy91) to sequentially integrate linear DNA cassettes containing OptoQAMP2-*PDC1*, mTagBFP2 and *Sp*Swi6/*Sp*Clr4, referred for simplicity as the *Sp*Swi6/*Sp*Clr4 strain. An isogenic control strain was made with the only difference being that it does not express *Sp*Swi6 and *Sp*Clr4. CENPK2.1C with or without integrations of *Sp*Swi6/*Sp*Clr4 was made to compare growth defect of the *Sp*Swi6/*Sp*Clr4 strain. See Extended Data Table 2 for strain genotypes, Extended Data Table 3 for plasmids used to generate the strains, and Extended Data Fig. 8 for plasmid maps.

### Chromatin compaction analysis

To analyze chromatin compaction, cells were stained with 4′,6-diamidino-2-phenylindole (DAPI) and values of fluorescent intensities were compared among strains. Briefly, cells were grown overnight and the next day, fresh media was inoculated with the overnight cultures and cells were incubated for 4 h. Cells were immobilized in Concanavalin A-coated well slides and fixation was performed by adding 4% paraformaldehyde (EMS 15714S), prepared in phosphate-buffered saline (PBS), and incubating 10 min. Next, cells were permeabilized with 0.2 % Triton X-100 in PBS, washed three times with PBS, and incubated with a DAPI aqueous solution 2 µg/ml (Sigma, D8417) for 5 min. After removing the DAPI solution and washing the cells with PBS, mounting media was added to the well slides before imaging. Microscopy was performed on a Zeiss Elyra PS.1 system. Structured illumination microscopy (SIM) images were acquired with a 63 × 1.46 NA oil objective in the DAPI channel with an exposure of 100 ms and 10 % laser power (4.1 mW 405 nm HR diode laser). Each image was acquired using 3 rotations and 5 phases per rotation. Images were processed using Zen Black structured illumination reconstruction algorithm.

### Nuclear morphology assessment

To visualize the nuclear envelope and nucleolus morphology, reference and mutant strains expressing fluorescent protein-fused marker proteins for both landmarks in the nucleus, Nup49-GFP and Nop1-mCherry, were grown overnight. These cultures were used to inoculate fresh media and cells were incubated until late logarithmic phase. Cells were then immobilized on ConA-coated well slides, and imaging was performed in a Zeiss Axio-Observer Z1 Yokogawa spinning disk confocal microscope using a 100 × 1.43 NA oil objective. For this, 10 Z-stack slices were recorded at a spacing of 300 nm and exposure times 50 ms and 100 ms, for GFP (50 %, 3 mW 488 nm excitation) and mCherry (50%, 3.3 mW 561 nm excitation) channels, respectively. Images were analyzed with Fiji software^112^.

### *INO1* locus statistical distribution

The procedure for the statistical mapping of the *INO1* locus was similar to previously described by Brickner *et al*.^110^ with slight modifications. In this case, the reference and *Sp*Swi6/*Sp*Clr4-expressing mutant strains with the *INO1* tagged with the 128 LacO array and the same nuclear landmark protein-fluorescent protein fusions described above were grown overnight in the corresponding dropout media without inositol or containing freshly prepared 100 µM *myo*-inositol. The next day, these cultures were diluted in fresh medium and incubated for approximately two generations. Cells were then immobilized in ConA-coated well slides and microscopy was performed as described for the nuclear morphology analysis but in this case 32 Z-stack slices were recorded for each field at a spacing of 250 nm. Nucloc software was used to process images in a modified mode where probability maps are presented as percentiles using a kernel density estimate^42^.

### *INO1* tracking and mechanical properties

To track the *INO1* gene, the same strains and culture conditions used for the statistical distribution of the locus analysis were reproduced. Cells were also immobilized on ConA-coated wells but in this case, time-lapsed images of each field in the GFP channel (100 mW 488 nm excitation) were recorded at intervals of 500 ms for 1 min. Fixed cells were also imaged with the same settings as controls for microscope stage drift. The WaveTracer tool on the Metamorph software was used to track the locus for each cell in each time-lapse set of images^113^. This tool enabled automatic segmentation and positioning of the loci in each slide to determine Mean Squared displacement (MSD) values for each locus as well as discard curves that do not have linear slopes.

Two mechanical properties of the *INO1* locus were calculated by using these MSD values and tracking coordinates. First, the radius of confinement (Rc) was calculated from the MSD plateaus (equation 1)^46,114^.

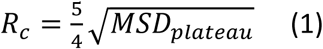

Second, the effective spring constant (*k_s_*) for the locus was calculated by quantifying the standard deviation (σ) of each step from the mean position by applying the equipartition theorem (Equation 2)^47,115^.

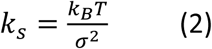

Student’s t-tests analyses were performed between the different strains and conditions to determine statistical differences for both *Rc* and *k_s_* parameters in each case.

### Mutation frequency measurements

#### Canr mutator assay

To measure the mutation frequency for a DBY746 *Sp*Swi6/*Sp*Cl4 expressing strain (ML35, Extended Data Table 2) versus reference (DBY746, Extended Data Table 2), we quantified the frequency of mutation of the *CAN1* (YEL063) gene. For this, both reference and *Sp*Swi6/*Sp*Clr4 mutant strains were incubated overnight and inoculated in 50 mL SC lacking methionine (SC-met). Cells were incubated over 13 days at 30°C and 200 rpm. To measure cell viability, every two days, samples of each culture were diluted to a final concentration of 10^3^ cells/ml and 100 μl of this suspension was inoculated on solid Yeast Extract–Peptone–Dextrose (YPD) medium. Colony-forming units (CFUs) were counted after 48 h. To quantify canavanine-resistant mutants (Can^r^), every two days, approximately 2 x 10^7^ cells were taken from each culture, washed with sterile H_2_O, and plated in SC lacking methionine and arginine (SC-met-arg) supplemented with 60 μg/ml L-canavanine sulfate. In this case colonies were counted after 3 to 4 days of incubation. To calculate frequency of mutants, the ratio of Can^r^/total viable cells was determined.

#### CAN1 sequencing

Sequencing was performed for 10 clones of the Can^r^ mutants obtained for each DBY746 and DBY746-*Sp*Swi6-*Sp*Cl4 strains after 9 days incubation in canavanine plates. Mutant colonies were collected, and genomic DNA was isolated by using a simple lithium acetate (LiOAc)-SDS procedure previously described^116^. A set of primers was used to amplify a region of 2082 bp including the whole *CAN1* open reading frame: CAN1-5UTR-Fw (5’-CAGAGTAAACCGAATCAGGGAATCCC-3’) and CAN1-3UTR-Rev (5’-GCTCATTGATCCCTTAAACTTTCTTTTCGG-3’). PCR products were purified and sent for sequencing using the amplification primers plus additional ones to completely cover the 2 kb fragment: CAN1-296-Fw (5’-AGACATATTGGTATGATTGCCCTTGG-3’), CAN1-562-Fw (5’-ATCACTTTTGCCCTGGAACTTAGTGTAG-3’), CAN1-989-Fw (5’-GAGCCATCAAAAAAGTTGTTTTCCGTATCTTAAC-3’).

#### Frequency of gross chromosomal rearrangement (GCRs)

To quantify GCRs, cultures with the DBY746 reference and DBY746-*Sp*Swi6-*Sp*Cl4 mutant strains transformed with the *URA3* cassette that replaced the *HXT13* locus (ML36 and ML37 strains respectively, Extended Data Table 2), were prepared as described before for the Can^r^ mutator assay. Cell viability was also measured by plating cells dilutions in YPD plates. To detect large rearrangements in the region containing both *CAN1* and *URA3,* every 4 days, approximately 10^8^ cells from each culture were washed with sterile H_2_O and plated in SC-met-arg plates containing 60 ug/ml L-canavanine and 1 mg/ml 5-fluoroorotic acid (5FOA). Colonies were counted after 3 to 4 days of incubation and the frequency of GCR events was calculated in a similar way as done for the Can^r^ mutants.

To calculate statistical differences for mutation frequencies between the different strains and conditions Student’s t-tests analyses were performed.

#### Whole-genome sequencing analysis

To determine differences in mutation numbers and patterns in the genome of *S. cerevisiae* expressing *Sp*Swi6 and *Sp*Clr4 compared to reference, WGS was performed. In this case cultures of the DBY746 reference and DBY746-*Sp*Swi6-*Sp*Cl4 mutant strains were set as before for the Can^r^ mutator and GCRs assays. After 13 days incubation, approximately 102 cells from each culture were washed with sterile H_2_O and plated in SC-met plates. Genomic DNA for 10 colonies from each strain as well as for the original reference and DBY746-*Sp*Swi6-*Sp*Cl4 was extracted using Zymo Research Corporation YeaStar™ Genomic DNA Kit (Zymo Research, 11-323). Quality of the DNA was assessed by assessed by agarose gel electrophoresis and concentration measurement before sending the samples for further processing at the IRIC Genomic platform.

Whole genome DNA libraries were generated from 50ng of DNA using the Kapa Hyperprep DNA library kit (Roche). Library quality was verified on a Bioanalyzer (Agilent), quantified using qPCR and sequenced on a NextSeq500 MidOutput 150cycles flowcell, obtaining around 10M Paired-End reads per sample (PE75).

To analyze the demultiplexed reads, first, we aligned the fastq files to the S288C yeast genome using Bowtie2 (version 2.2.9) ^117^ and converted them to BAM files with SAMtools (version 1.3.1)^118^. During the alignment process, Bowtie2 was configured to allow a maximum of two distinct alignments per read, and only the sequencing reads that aligned in the SAM output were retained. Our alignment success rates ranged from 85% to 98%, Subsequently, we identified variants in the strains by utilizing the mpileup function of SAMtools ^118^, which utilized the output from the Bowtie2 alignment. The variant information was then saved in Variant Call Format (VCF)^119^. For variant calling, BCFtools (version 1.3.1)^120^ employed genotype likelihoods to identify SNPs and indels. To compare the mutations found in our mutant strains with those in the control strain, we filtered out the variants present in the mutant samples that were also present in the control samples, subsequently we incorporated the variants found in the control strains but absent in the mutant as these variants represented distinct mutations differentiating the mutant and control strains. Variants with a quality score greater than 20 and read depth higher than 50 were retained for the further analysis.

### Recombination rate measurements

#### Sporulation and flow cytometry procedure

Sporulation and flow cytometry analysis of the spores was performed as described before^61^ with some variations. Diploids strains were inoculated on 5 ml YPD and incubated overnight at 30°C, 200 rpm. Cells were harvested at 2000 rpm, 2 min and resuspended in 600 μl of sterile H_2_O, and 150 μl of each preparation was plated on four different plates containing sporulation medium (2% potassium acetate, 1.2 % agar, and 10 µg/ml of L adenine, L arginine, L histidine, L leucine, L lysine, L phenylalanine, L threonine, L tryptophan, L uracil, L valine). Plates were incubated for 10 days at 30°C. After this time, cells were recovered using polystyrene scraper loops and were resuspended in tubes containing 5 mg/ml zymolyase 100T (Zym002.250, Bioshop) in 750 μl of sterile H_2_O and 100 μl glass beads (Glass beads, acid-washed, 425-600 μm, G8772, Sigma). Tetrads were then disrupted by vortexing the tubes in a Mini-BeadBeater-16 (Biospec), for 1 min, followed by incubation at 30°C for 60 min, and another cycle of vortexing for 1 min. Samples were centrifuged 5 min at 4500 rpm and pellets were resuspended by vortexing in 200 μl of sterile H_2_O. Samples were centrifuged again and the supernatants containing mainly vegetative cells were discarded. Spores were resuspended by vortexing in 600 μl sterile H_2_O containing 0.01% nonidet NP40. Spores were analyzed on a Bio-Rad YETI (ZE5) Cell analyzer and the Everest 3.0 acquisition software. A gate was set on the FSC *versus* SSC plot to only allow the analysis of events corresponding to spores (Extended Data Fig. 9). Cerulean, Venus and mCherry fluorescent markers were excited using a 405 nm, 488 nm and 561 nm laser respectively and read through a 525/50, 549/15 and 615/24 bandpass filter (Extended Data Fig. 10). Data were then analyzed using FlowJo V10 software.

#### Analysis of fluorescent patterns

As described above^61^, each of the tester strains contain three fluorescent markers corresponding to cerulean, yellow, and red (C, Y or R) channels. Thus, diploids cells that were obtained by crosses with haploids not containing any fluorescent marker produced eight different classes of spores (Fig. 4a). When analyzing the data from flow cytometry, each class was denoted by stating the markers presence in a consecutive array, and in the absence of fluorescence for any marker the symbol ‘+’ was used. For example, if the arrangement of the markers in the tri-fluorescent haploid parental strain was CYR, then parental spores containing all three markers or none were indicated as CYR and +++, respectively. The rest of the classes corresponded to single recombination events between C and Y markers (C++ and +YR), Y and R markers (CY+ and ++R), or double recombination events (C+R and +Y+) (Fig. 4a). Three projections were used to represent the intensities of two different fluorescence markers (Fig. 4c) where the nomenclature [X/+] specifies that the X marker might be present or not. As done previously, to classify each spore in one of the eight different classes that could be obtained, the following rules to analyse the quadrants in each projection were applied:

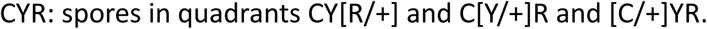

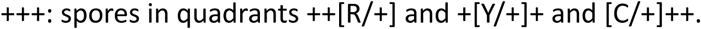

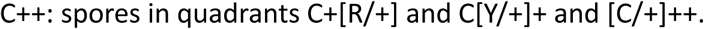

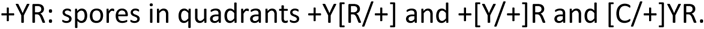

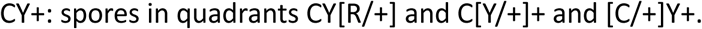

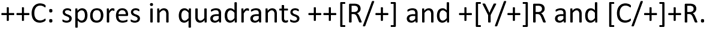

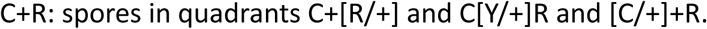

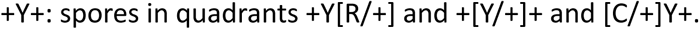

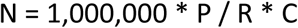

#### Recombination rate quantification

To calculate the recombination rates for two linked markers, for example C and Y, the following equation (3) was used:

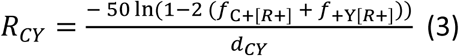

In this case *d*CY represents the distance in kbp between the markers, and *f*XYZ is the frequency of the particular XYZ class of spores. The Values for recombination rate were expressed in centiMorgan (cM) per kilobase.

### Co-immunoprecipitation

Co-IP experiments between *Sp*Swi6 and H3K9me2 or H3K9me3 in *Sp*Swi6/*Sp*Clr4 (no tag control), *Sp*Swi6-Flag/*Sp*Clr4 and SpSwi6-5Flag cells were performed. Yeast strains were grown in 50 mL of yeast nitrogen base (YNB) medium lacking methionine (YNB-Met) to an OD_600_ of 0.8, collected by centrifugation, washed in cold ddH_2_O, and resuspended in 700 μL cold IP buffer (20 mM HEPES-KOH pH 7.5, 150 mM NaCl, 1 mM EDTA pH 8.0, 5 mM MgCl_2_, 0.25% Triton X-100, 10% glycerol, 0.5 mM DTT and protein inhibitors (1 mM PMSF, 1 mM Benzamidine, 10 μg/mL Aprotinin, 1 μg/mL Leupeptin, 1 μg/mL Pepstatin)). About the same amount of OD_600_ units was used for all samples. Cells were lysed by bead beating for 5 min and the soluble extract was recovered by centrifugation (15 min at 4°C at 18,000 × g). About 3-3.5 mg of protein (600 µl) was taken per IP and 6 μL (1%) was saved as Input sample. Protein extracts were incubated for 90 min at 4°C with 50 μL of pre-washed Dynabeads coated with Pan Mouse IgG antibodies (Thermo Fisher Scientific, 11042) coupled to 5 μg of anti-Flag mouse monoclonal antibody (Sigma-Aldrich, F3165). Samples were washed three times with cold IP buffer and resuspended in Laemmli buffer. Samples were then incubated at 95°C for 5 min, briefly centrifuged, and analyzed by SDS-PAGE and western blot using anti-Flag mouse monoclonal (Sigma-Aldrich, F3165; 1:2,000 dilution), anti-H3K9me2 mouse monoclonal (Abcam, ab1220; 1:1,000 dilution) and H3K9me3 rabbit polyclonal (Abcam, ab8898; 1:1,000 dilution) antibodies. Membranes were incubated with donkey anti-mouse IRDye 680RD antibodies or donkey anti-rabbit IRDye 800CW (LI-COR Biosciences, 926-32213 and 926-68072) according to the manufacturers’ instructions and scanned on the Odyssey infrared imaging system (LI-COR Biosciences).

### ChIP-seq

Yeast ChIP experiments were performed from two independent biological replicates according to^121^. In brief, yeast cultures were grown in 50 mL of YNB-Met medium to an OD_600_ of 0.7-0.9 before crosslinking with 1% formaldehyde (Fisher Scientific, BP531-500) at room temperature for 30 min and quenched with 125 mM glycine. Crosslinked cells were collected by centrifugation and washed twice with 1X TBS (20 mM Tris-HCl pH 7.5, 150 mM NaCl). Cell pellets were then resuspended in 700 μL Lysis buffer (50 mM HEPES-KOH pH 7.5, 140 mM NaCl, 1 mM EDTA, 1% Triton X-100, 0.1% Na-deoxycholate and protease inhibitor cocktail (1 mM PMSF, 1 mM Benzamidine, 10 μg/mL Aprotinin, 1 μg/mL Leupeptin, 1 μg/mL Pepstatin). About the same number of OD_600_ units was used for all samples. A fixed amount of crosslinked *S. pombe* cells expressing 3Flag-Swi6 (a kind gift from D. Moazed), representing 10% of the *S. cerevisiae* cells, were added to the *S. cerevisiae* cells before cell lysis. This was used to normalise samples, hence allowing sample-to-sample comparisons, and validating the ChIP. Cells were lysed by bead beating and the lysate was sonicated with a Model 100 Sonic dismembrator equipped with a microprobe (Fisher Scientific), 4 x 20 sec at output 7 Watts, with a 1 min break between sonication cycles. Soluble fragmented chromatin was recovered by centrifugation. 600 μL of the chromatin sample was taken per immunoprecipitation (IP) and 6 μL (1%) was saved as an Input sample. The following amounts of antibody per IP were used: anti-Flag (Sigma-Aldrich, F3165; 5 μg), anti-H3 (Abcam, ab1791; 2 μg) and anti-H3K9me2 (Abcam, ab1220; 2 μg). All antibodies have been validated for ChIP (see manufacturers’ websites). Anti-Flag and anti-H3K9me2 antibodies were coupled to Dynabeads coated with Pan Mouse IgG antibodies (Thermo Fisher Scientific, 11042) and anti-H3 antibody was coupled to a mix of Dynabeads coated with Protein A (Thermo Fisher Scientific, 10002D) and Dynabeads coated with Protein G (Thermo Fisher Scientific, 10004D). 50 μL of the appropriate Dynabeads pre-coupled with the indicated antibody were added to the chromatin sample and incubated overnight at 4°C. Beads were washed twice with Lysis buffer, twice with Lysis buffer 500 (Lysis buffer + 360 mM NaCl), twice with Wash buffer (10 mM Tris-HCl pH 8.0, 250 mM LiCl, 0.5% NP40, 0.5% Na-deoxycholate, 1 mM EDTA) and once with TE (10 mM Tris-HCl pH 8.0, 1 mM EDTA). Input and immunoprecipitated chromatin were eluted and reverse-crosslinked with 50 μL TE/SDS (TE + 1% SDS) by incubating overnight at 65°C. Samples were treated with RNase A (345 μL TE, 3 μL 10 mg/mL RNAse A (Sigma-Aldrich, R6513), 2 μL 20 mg/mL Glycogen (Roche, 10901393001)) at 37°C for 2 hr and subsequently subjected to Proteinase K (15 μL 10% SDS, 7.5 μL 20 mg/mL Proteinase K (Thermo Fisher Scientific, 25530049)) digestion at 37°C for 2 hr. Samples were extracted twice with phenol/chloroform/isoamyl alcohol (25:24:1), followed by precipitation with 200 mM NaCl and 70% ethanol. Precipitated DNA was resuspended in 50 μL of TE before being used in ChIP-seq experiments.

For the Flag ChIPs, to obtain enough material for the library preparation, four technical replicates were pooled. All ChIP and Input samples (see above) were subjected to a 1.7X cleanup using KAPA Pure Beads (Roche, 07983280001) according to the manufacturers’ instructions. Samples were eluted with 40 μL of Elution buffer (10 mM Tris pH 8.0). DNA concentration of ChIP and Input samples was determined by qPCR using a standard curve made with a fragment size control sample. 1-10 ng of ChIP and Input DNA were used for ChIP-seq library preparation as follow. The ends of DNA were repaired by incubating in 70 μL of 1X NEBuffer 2 containing 0.6 units of T4 DNA polymerase, 2 units of T4 polynucleotide kinase (NEB, M0201S), 0.09 nM dNTPs and 0.045 μg/μL of BSA at 12°C for 30 min. Repaired DNA was then subjected to a 1.7X cleanup using KAPA Pure Beads before dA tailing as follow. Beads containing the repaired DNA were resuspended in 50 μL of 1X NEBuffer 2 containing 0.1 mM dATP and 25 units of Klenow Fragment (3’→5’ exo-) (NEB, M0212M), and incubated at 37°C for 30 min. After a 1.8X cleanup with KAPA Pure Beads, the A-tailed DNA was ligated to index adapters (Illumina Truseq DNA UD Indexes (20023784)) as follow. The beads were resuspended in 45 μL of 1X Ligase buffer containing 8 nM of adapter and 2.5 units of T4 DNA ligase and incubated at room temperature for 60 min. The ligated DNA was then subjected to a 1X cleanup with KAPA Pure Beads, followed by a double size selection (0.52X-1X) leading to fragments in the 200-600 bp range. Libraries were PCR-amplified with 11-12 cycles using KOD Hot Start DNA polymerase (Millipore, 71086-3) and cleaned up using 1X KAPA Pure Beads. Libraries were qualified on Agilent 2100 Bioanalyzer using High Sensitivity DNA Kit and quantified by qPCR using NEBNext Library Quant Kit for Illumina (NEB, E7630). The equal molarity of each library was pooled and subjected to sequencing on Illumina NextSeq 500 platform at the Institute for Research in Immunology and Cancer (IRIC) to generate 75 bp pair-end reads.

### ChIP-seq Data analysis

Adapter sequences were removed from paired-end reads for each biological replicate using Trimmomatic (version 0.36)^122^ with parameters “ILLUMINACLIP:adapters.fa:2:30:10: TRAILING:3 MINLEN:25“. Paired-end reads for each biological replicate were independently aligned to the S. cerevisiae (UCSC sacCer3) and S. pombe (Downloaded from https://www.pombase.org/ on December 7th 2022) reference genomes using the short read aligner Bowtie 2 (version 2.3.4.3)^123^. Only reads mapped in proper pairs and primary alignments were kept in aligned files using Samtools (version 1.9)^124^ with parameters “-F 2048 -F 256 -f 2”. Duplicate reads were removed from aligned files using Samtools “fixmate” and “markdup -r”. Coverage for each base pair of the S. cerevisiae genome was computed using genomeCoverageBed from BEDTools (version 2.27.1)^125^ and normalized using S. pombe reads as follows. The read density at each position of the S. cerevisiae genome was multiplied by a normalization factor N defined as:

N = 1,000,000 * P / R * C
where:

1,000,000 is an arbitrary chosen number used for convenience; R is the total number of reads in the IP sample that mapped to the *S. pombe* genome; C is the total number of reads in the Input sample that mapped to the *S. cerevisiae* genome; P is the total number of reads in the Input sample that mapped to the S. pombe genome.

The genome coverage files were converted to the bigWig format using the utilities from UCSC Genome Browser^126^.

Correlation plots were generated using deepTools (version 3.5.1)^127^ with parameters “multiBigwigSummary bins -- binSize 100” and “plotCorrelation -- corMethod pearson -- whatToPlot heatmap --colorMap PiYG --plotNumbers”.

Statistics regarding the reads (including the number of reads, mapped reads, the correlation between replicates) are shown in Supplementary Table 1. Note that replicate 2 from the H3K9me2 ChIP experiment from *Sp*Swi6-Flag/*Sp*Clr4 cells failed and was omitted from our analyses.

Coverage files (BigWig) files were visualized on the UCSC Genome Browser^126^.

Aggregate profiles (metagene analyses) were generated using the Versatile Aggregate Profiler (VAP)^128,129^. The aggregate profiles were generated by aligning data on the transcription start site (TSS) of all verified genes, excluding those with signal >5 rpm in the no tag control samples. The data was averaged over 10 bp bins. TSS coordinates were from^130^. The metagene includes 3896 genes, listed in Supplementary Table 2.

### Fermentation experiments and data analysis

Yeast strains were streaked from a −80 °C glycerol stock to yeast extract peptone agar plates supplemented with 3% v/v glycerol and 2% v/v ethanol and incubated in dark at 30 °C. One colony was selected to grow overnight in SC-GE media in blue light at 30 °C. After overnight growth, cells were diluted to an OD_600_ of 0.01 in 20 mL of minimal media^131^, containing per liter: 100 mg calcium chloride, 100 mg sodium chloride, 500 mg magnesium sulfate, 5 g ammonium sulfate, 1 g potassium chloride, 500 µg boric acid, 40 µg copper sulfate, 100 µg potassium iodide, 200 µg ferric chloride, 400 µg manganese sulfate, 200 µg sodium molybdate, 400 µg zinc sulfate, 1 µg biotin, 200 µg calcium pantothenate, 1 mg inositol, 200 µg niacin, 100 µg p-aminobenzoic acid, 200 µg pyridoxine, 200 µg thiamine, 10 mg potassium phosphate and 20 g glucose. A volume of 20 mL of diluted cells (OD_600_ = 0.01) were dispensed in sterile 30 mL glass vials and placed in ChiBio mini bioreactors^83^. For repression assays, doxycycline and methionine were added to final concentrations of 2 μg/mL and 1mM respectively. Fermentations were incubated at 30 °C with stirring set to 0.5. When required, the 457 nm LED was used to expose cultures to blue light with a 0.1 intensity setting and on/off frequency of 10/100 s. Turbidity readings were automatically taken by the mini bioreactor units every minute. After the end of fermentations, samples of cheater strains were taken from reactors grown in dark to make glycerol stocks and screened for BFP fluorescence by flow cytometry (Beckman and Coulter Cytoflex) with a 405 nm excitation laser and 450/45 bandpass, to ensure growth of samples was not due to contamination (Extended Data Fig. 11).

ChiBio turbidity readings were converted to absorbance units at λ 600 nm with a calibration curve (Extended Data Fig. 12). For data visualization, data was smoothed by moving average (n=5). Culture stability was estimated by measuring the duration of the lag phase using the tangent method^13^, in what should have been non-permissive conditions (darkness). To calculate the lag phase, we used the first observed maximum growth rate and a local regression of 9 points as parameters (Representative plots are shown in Extended Data Fig. 13). An unequal variance two-sided t test (Welch’s test, α=0.05) was used to determine statistical significance.

## Data availability

All raw and processed ChIP-seq data generated in this study have been submitted to the NCBI Gene Expression Omnibus (GEO; http://www.ncbi.nlm.nih.gov/geo/) under accession number GSE239679.

To review GEO accession GSE239679:

Go to https://www.ncbi.nlm.nih.gov/geo/query/acc.cgi?acc=GSE239679 Enter token ulajauwolfgjlmv into the box

## Extended Data

**Extended Data Fig. 1.**
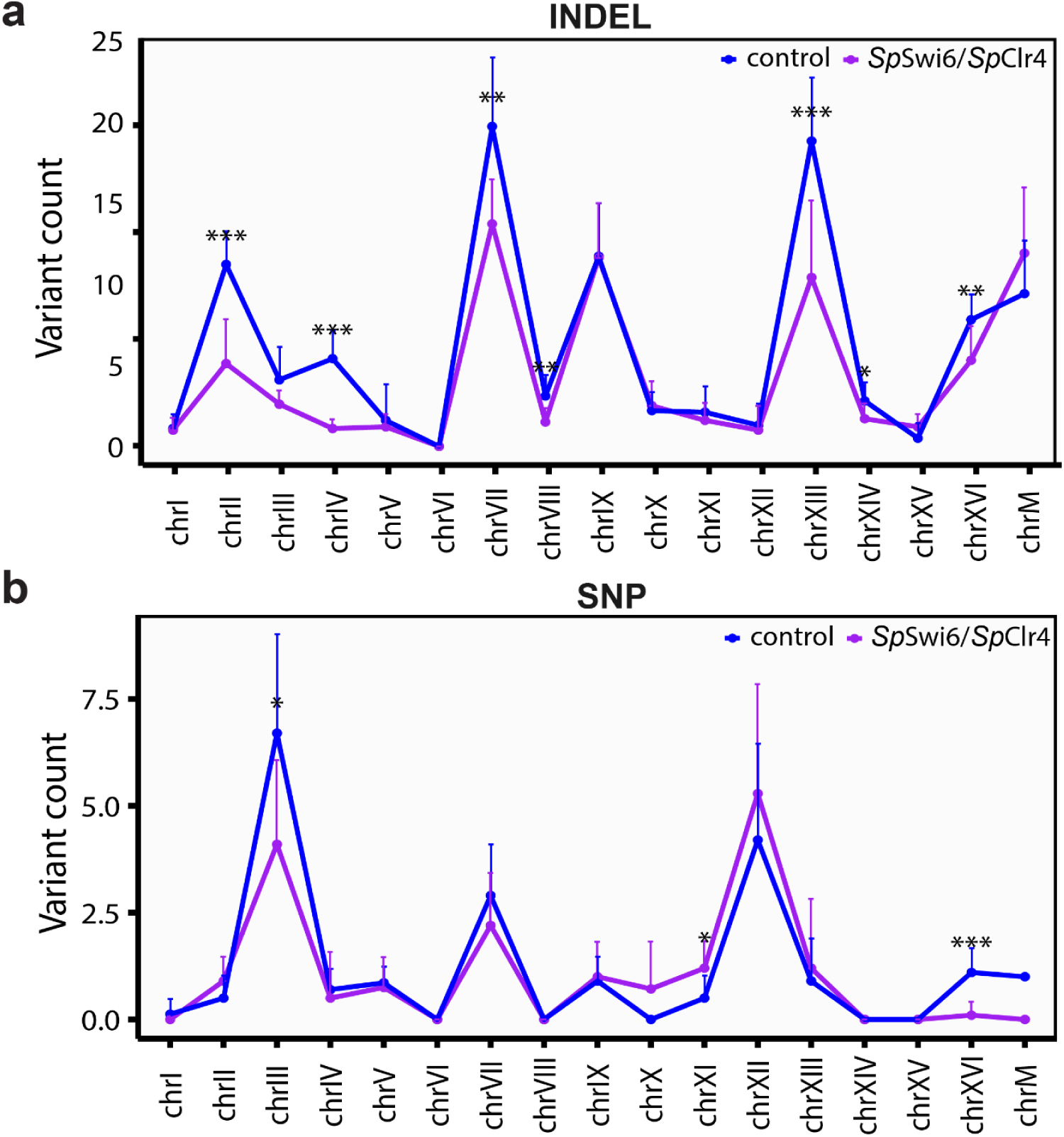
Variant count per chromosome in reference (DBY746) versus the DBY746-SpSwi6-SpClr4 strain after 13 days incubation for indel (**a**) and SNPs (**b**) mutations respectively.

**Extended Data Fig. 2.**
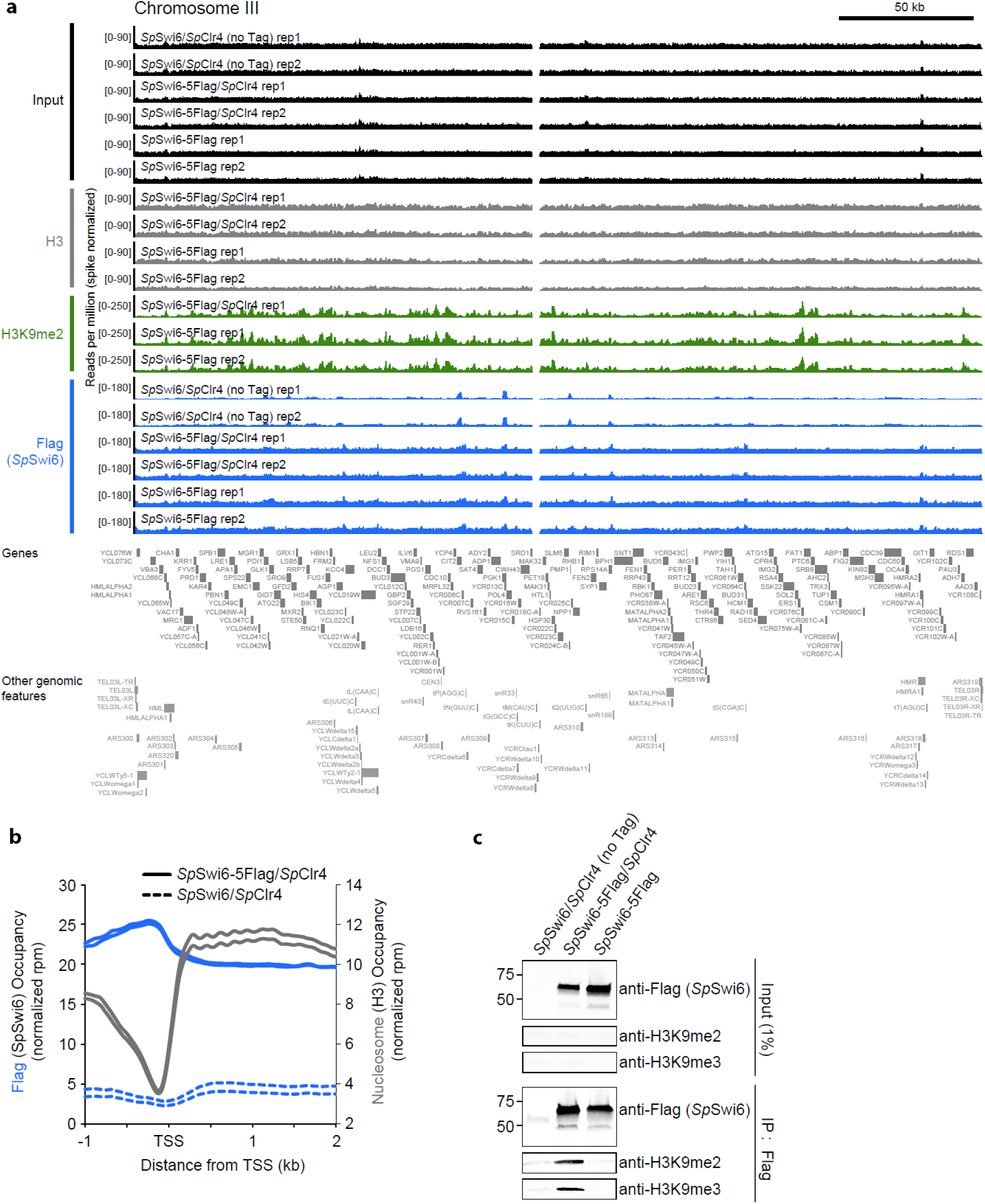
Genomic occupancy of H3, H3K9me2 and SpSwi6 in S. cerevisiae cells expressing SpClr4 and/or SpSwi6. **a,** A genome browser view of Inputs (black), H3 (grey), H3K9me2 (green) and anti-Flag (SpSwi6; blue)) ChIP-seq over the S. cerevisiae chromosome III. S. pombe-spike normalized read counts are shown for both replicates from ChIPs performed in S. cerevisiae cells expressing SpSwi6-5Flag/SpClr4 or SpSwi6-5Flag (no SpClr4 control). Data from SpSwi6/SpClr4 (no tag control) is also shown for the Inputs and the anti-Flag ChIP samples. Genes and other genomic features are shown below the data tracks. **b,** A metagene of H3 occupancy from SpSwi6-5Flag/SpClr4-expressing S. cerevisiae cells (grey) together with SpSwi6-5Flag occupancy from SpSwi6-5Flag/SpClr4-expressing (solid blue) and SpSwi6/SpClr4-expressing (no tag control; dashed blue) S. cerevisiae cells. **c,** Co-IP experiments between SpSwi6 and H3K9me2 or H3K9me3 in SpSwi6/SpClr4 (no tag control), SpSwi6-Flag/SpClr4 and SpSwi6-5Flag cells.

**Extended Data Fig. 3.**
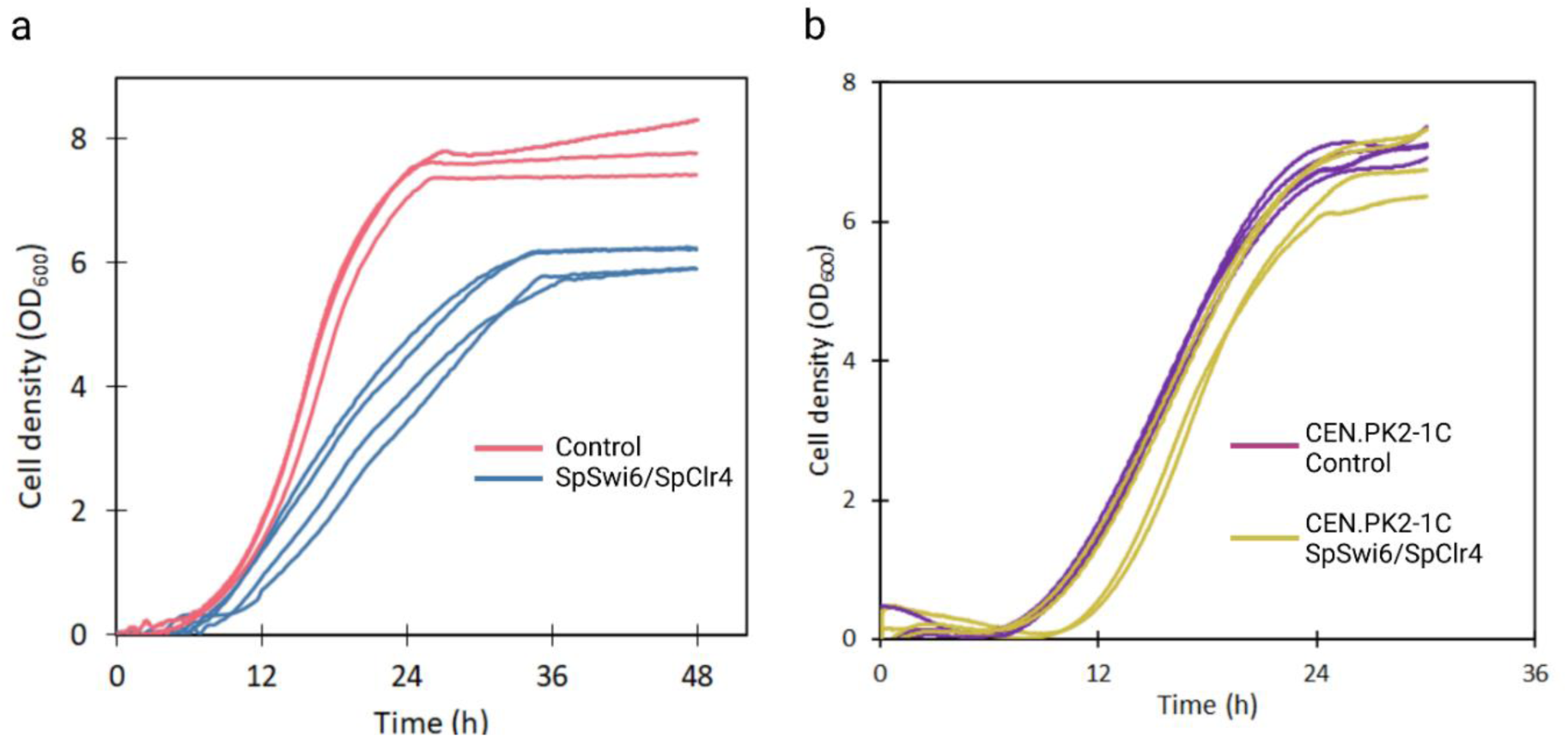
Effect of expressing SpSwi6/SpClr4 on cell growth. **a,** Growth curves of optogenetically controlled strains expressing SpSwi6/SpClr4 SGy162, blue, n=4) and control strain without SpSwi6/SpClr4 (SGY139, red, n=3) in glucose media and blue light (permissive conditions). SGy139 control strain grows faster and further than SGy162, which expresses SpSwi6 and SpClr4. **b,** Growth curves of reference strain (CEN.PK2-1C) expressing SpSwi6/SpClr4 (SGy164, green, n=4) or control strain without SpSwi6/SpClr4 (SGy163, yellow, n=4).

**Extended Data Fig. 4.**
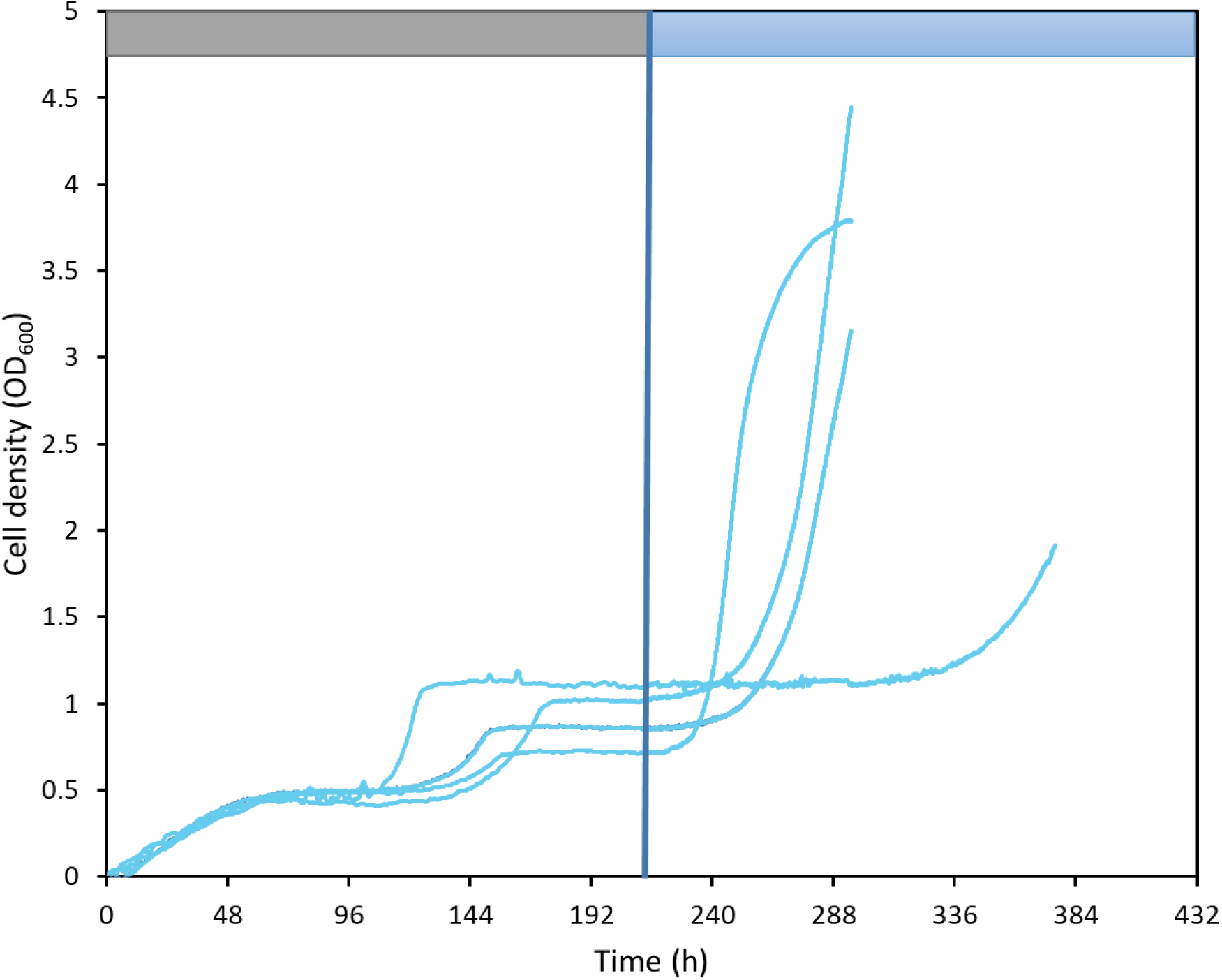
Growth curves of optogenetic strain replica cultures expressing SpSwi6/SpClr4 that never cheated during prolonged non-permissive dark incubations (Fig. 1c). When cultures were switched to blue light illumination conditions (vertical blue line, at t = 217 h), all culture were able to grow. Top panel depicts the periods when the cultures were exposed to darkness (left, gray) and light (right, blue). The fact that all cultures were able to grow when switched to light conditions, confirms that cells were alive during the prolonged repression period and remained stably light-regulated for the duration of the experiment.

**Extended Data Fig. 5.**
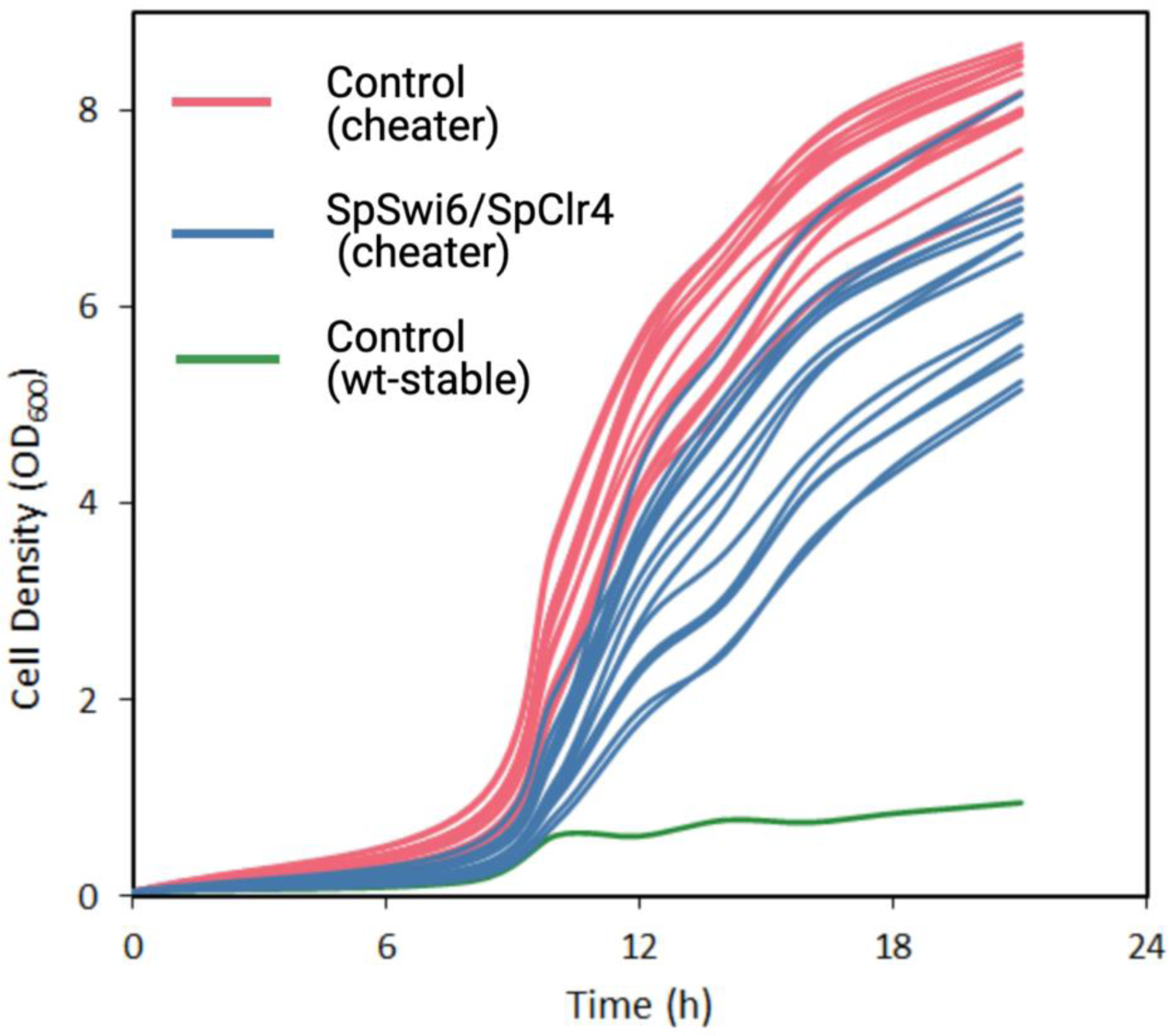
Growth curves of cheater cells derived from experiments in Fig. 1c (red and blue) compared to a stable optogenetically controlled strain (green). These plots reveal a complete loss of light regulation of cell growth in all cheater cells, indicative of mutation.

**Extended Data Fig. 6.**
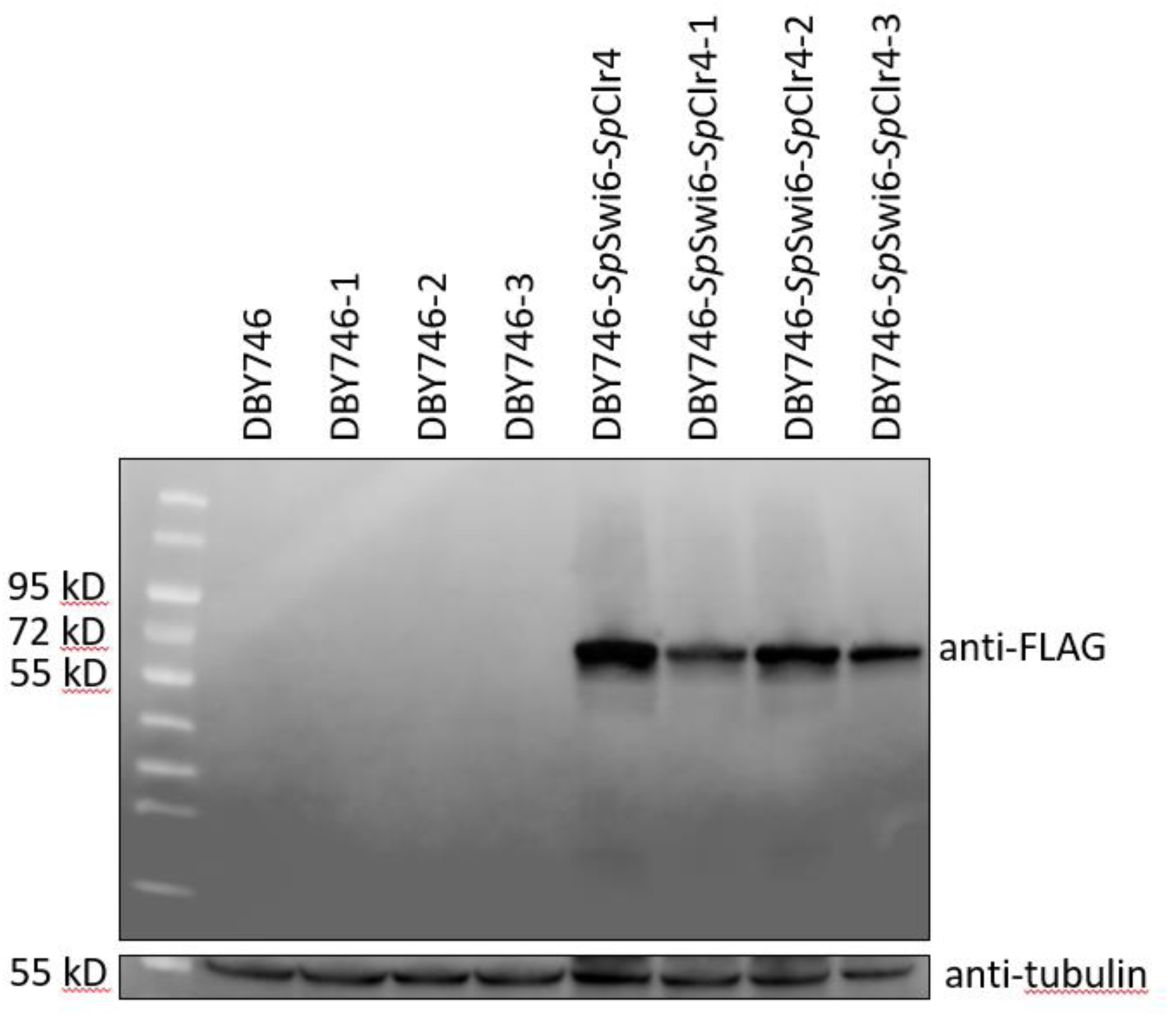
Detection of SpSwi6-5Flag in DBY746 strain and mutants. Protein levels in lysates from DBY746 strains used for the mutation frequency assays, carrying or not SpSwi6 tagged at C-terminal position with an array of 5x FLAG epitopes. Membranes were analyzed by western blotting with an anti-Flag M2 antibody.

**Extended Data Fig. 7.**
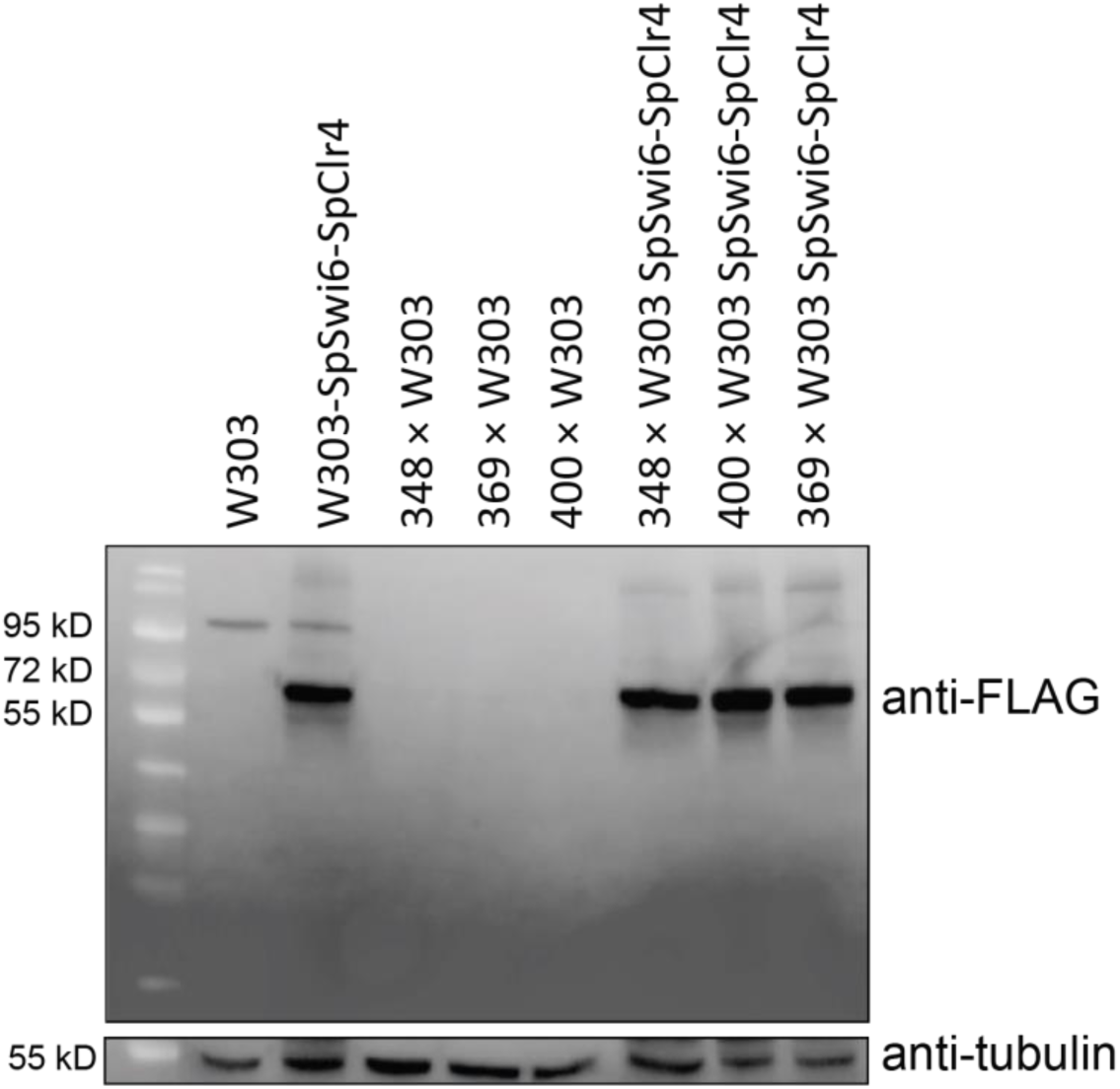
Detection of SpSwi6-5Flag in tri-fluorescent tester strains and mutants. Protein levels in cell extracts from different strains used for the recombination rate assays, expressing or not SpSwi6 fused to an array of 5x Flag epitopes at C-terminal position, and analyzed by western blotting with an anti-Flag M2 antibody.

**Extended Data Fig. 8.**
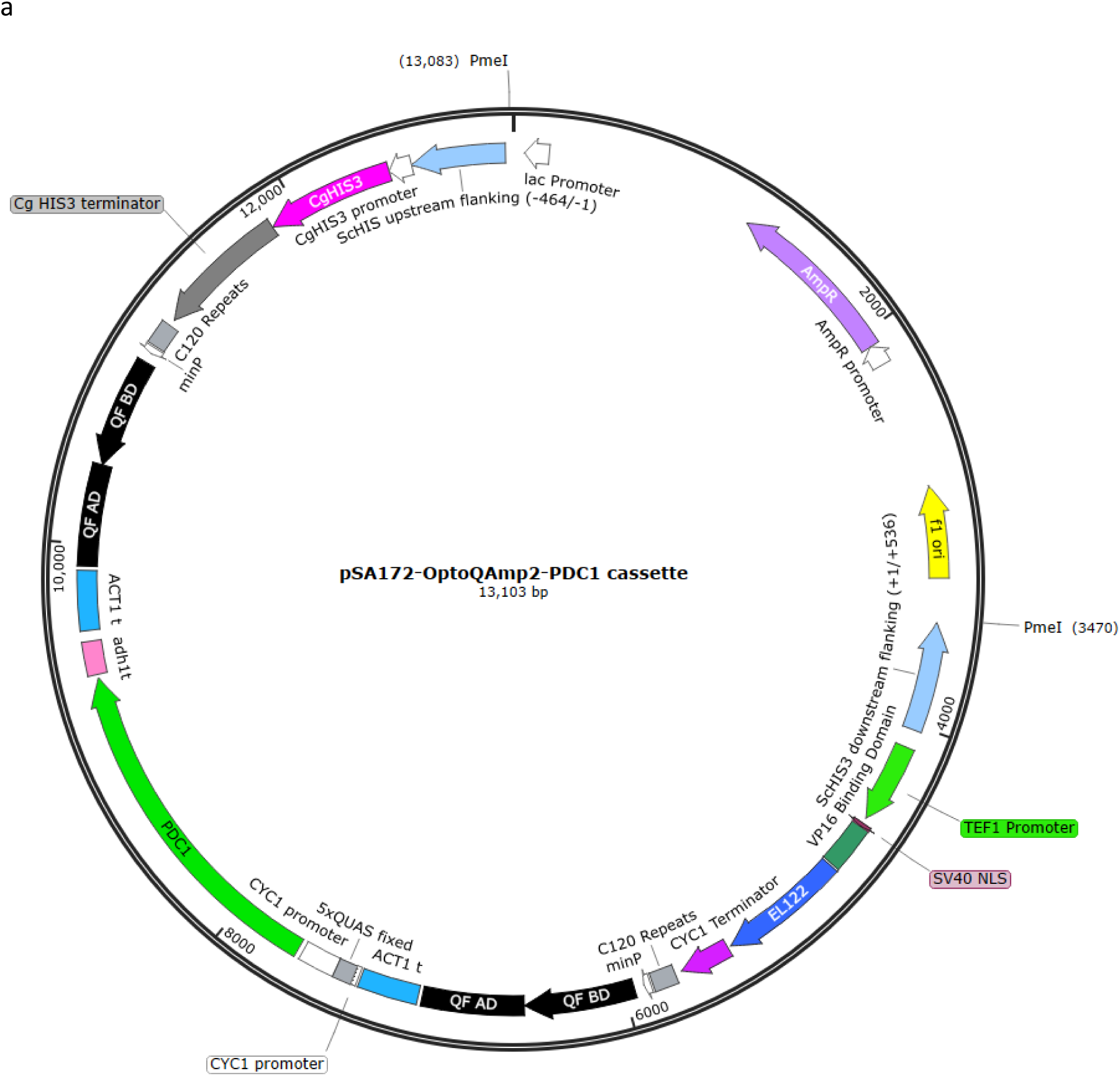
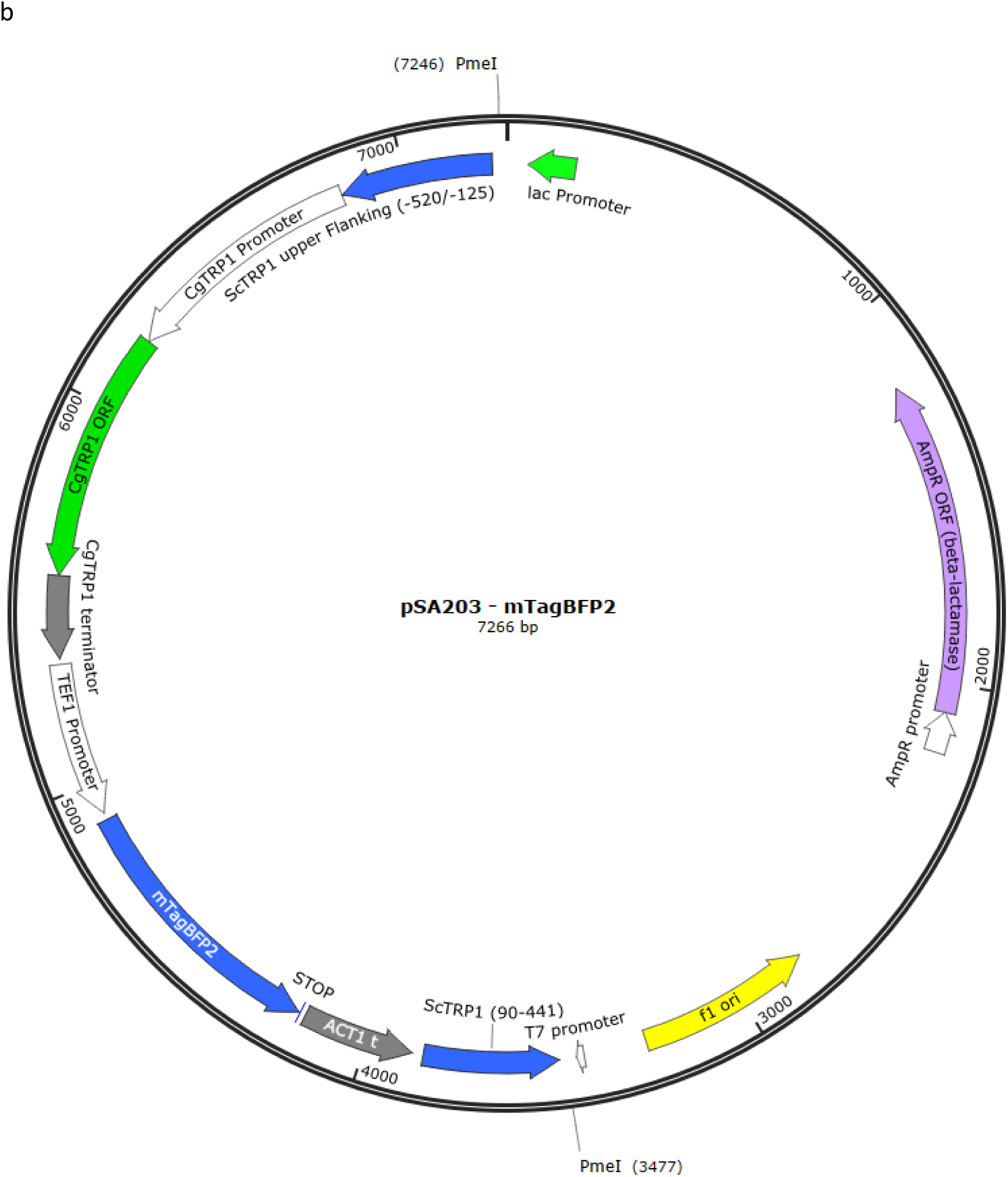
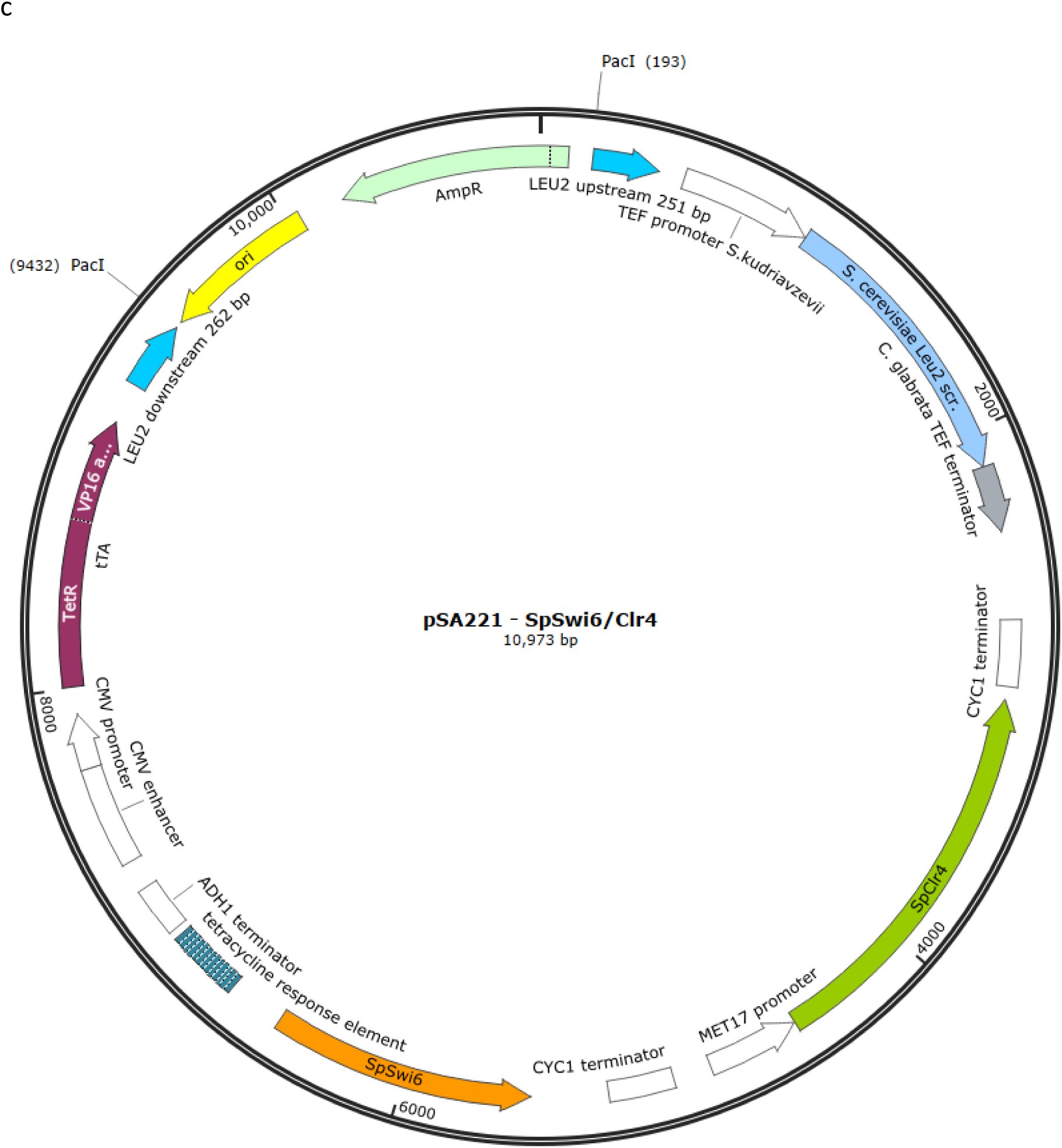
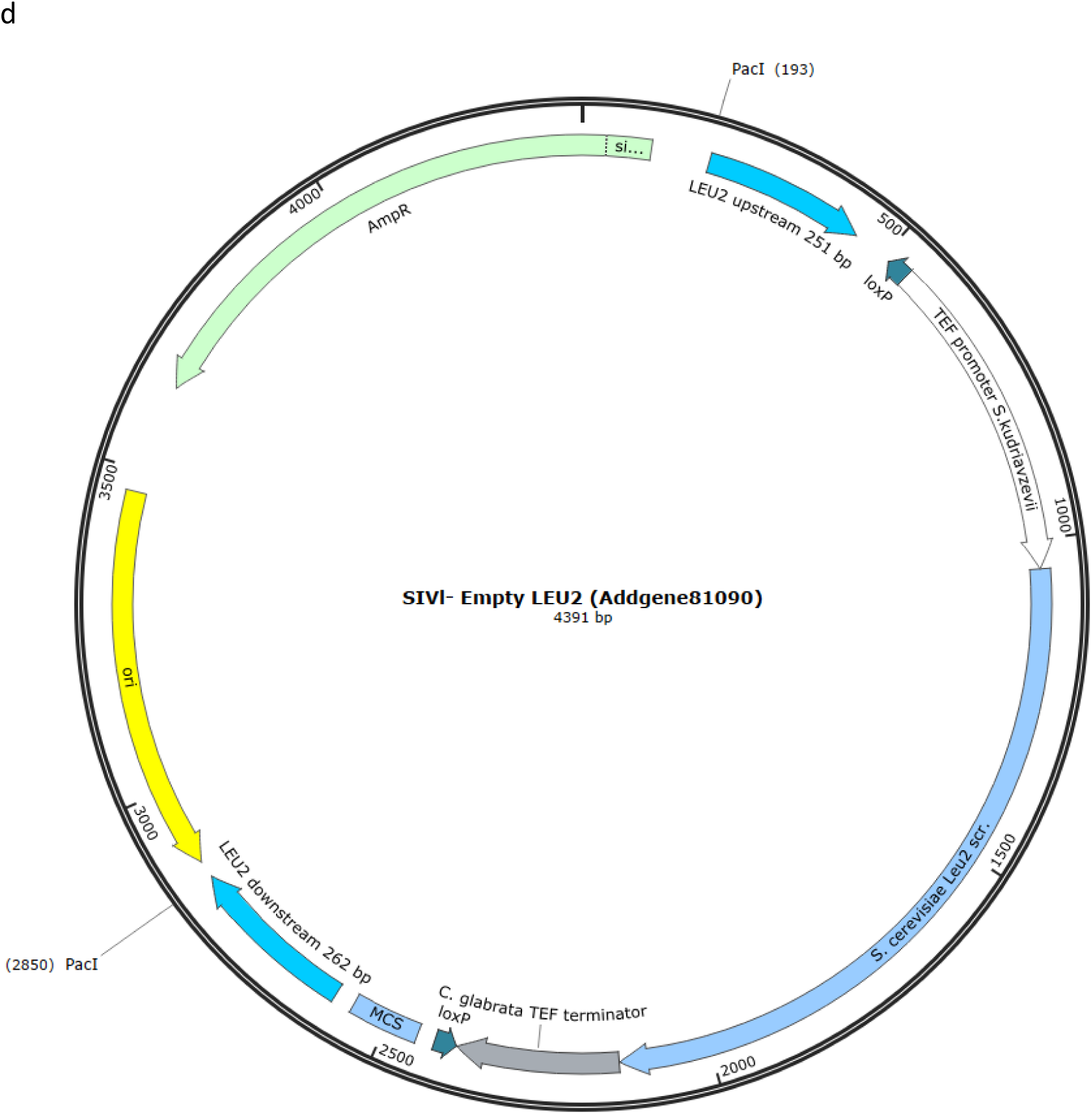
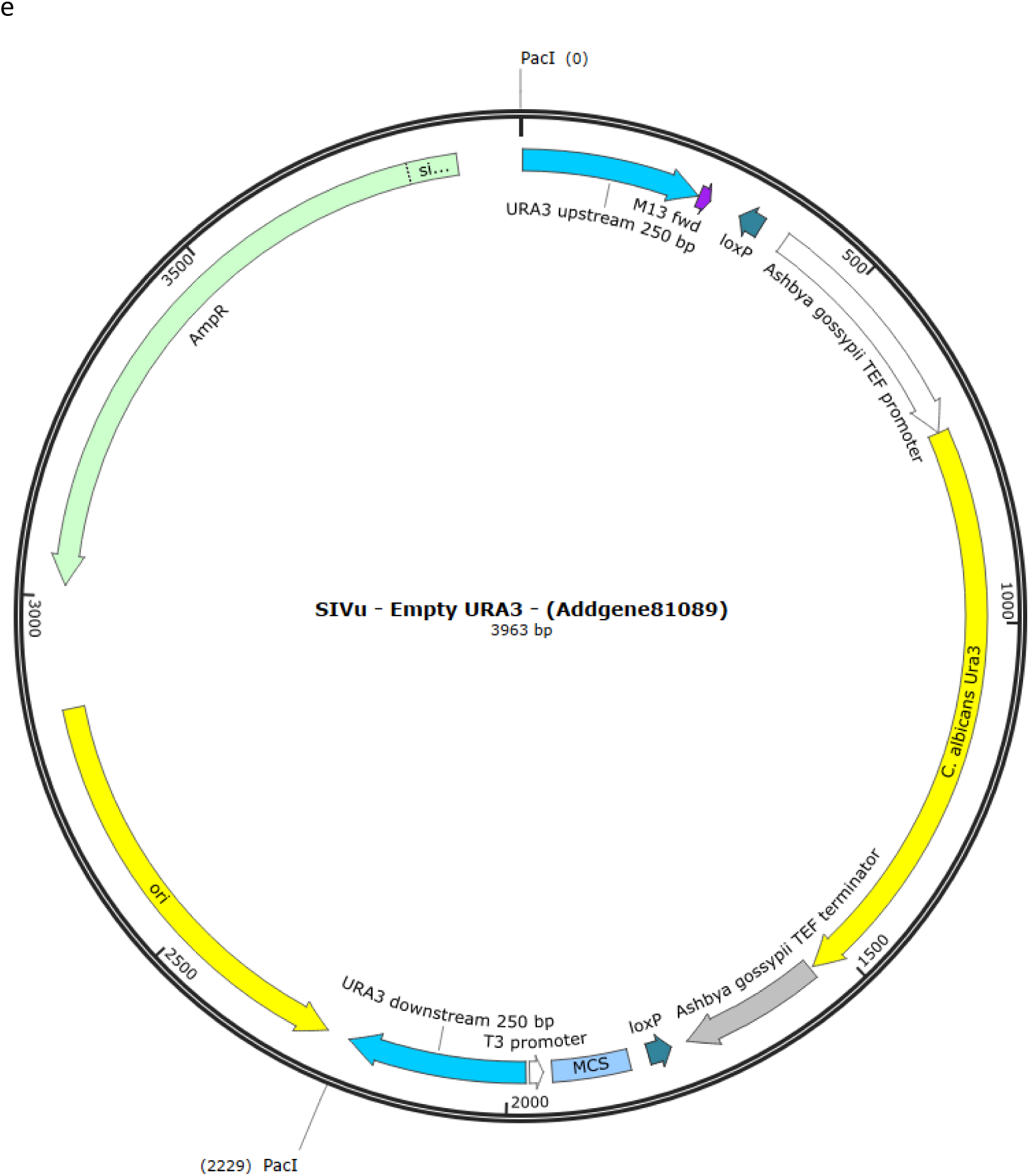
Maps of key plasmids used in this study. **a,** pSA172, **b,** pSA203, **c,** pSA221, Addgene81089, **d,** SIVl Addgene81090, **e,** SIVu. See Extended Data Table 3.

**Extended Data Fig. 9.**
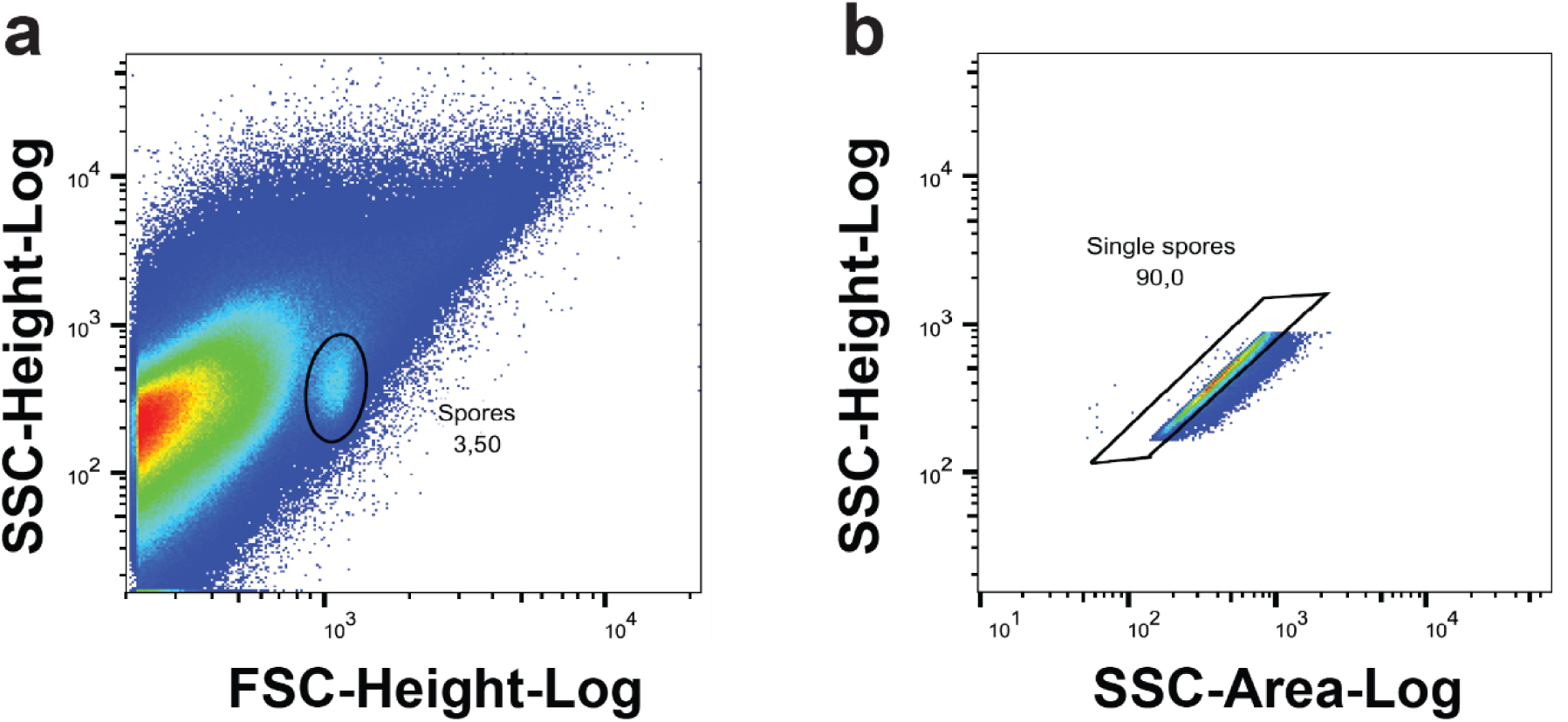
Spore events selection based on scattered light. **a**, Spores selection by using gates in the SSC-Height-Log versus FSC-Height-Log graph based on their size. **b**, Selection of single spores by using gates in the SSC-Height-Log versus SSC-Area-Log graph since events that correspond to doublets generate a lower ratio height vs area. Warm colors indicate highest density.

**Extended Data Fig. 10.**
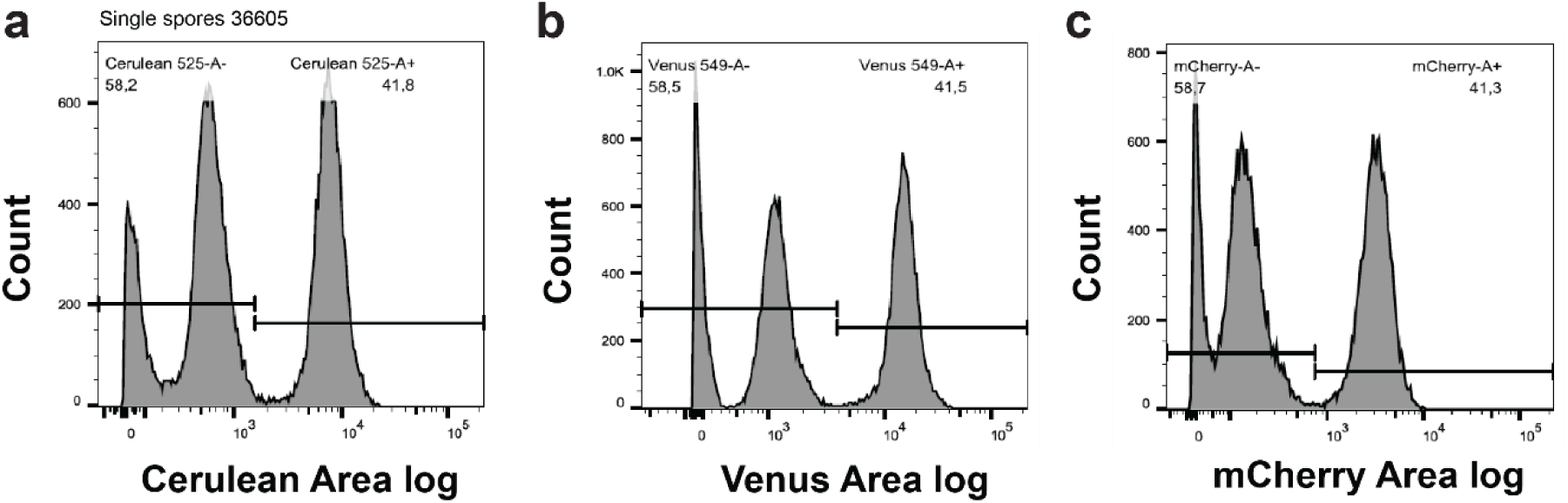
Fluorescence intensity distribution in spores. Example of spores from a 348 × W303 diploid hemizygote for markers C, Y, and R.

**Extended Data Fig. 11.**
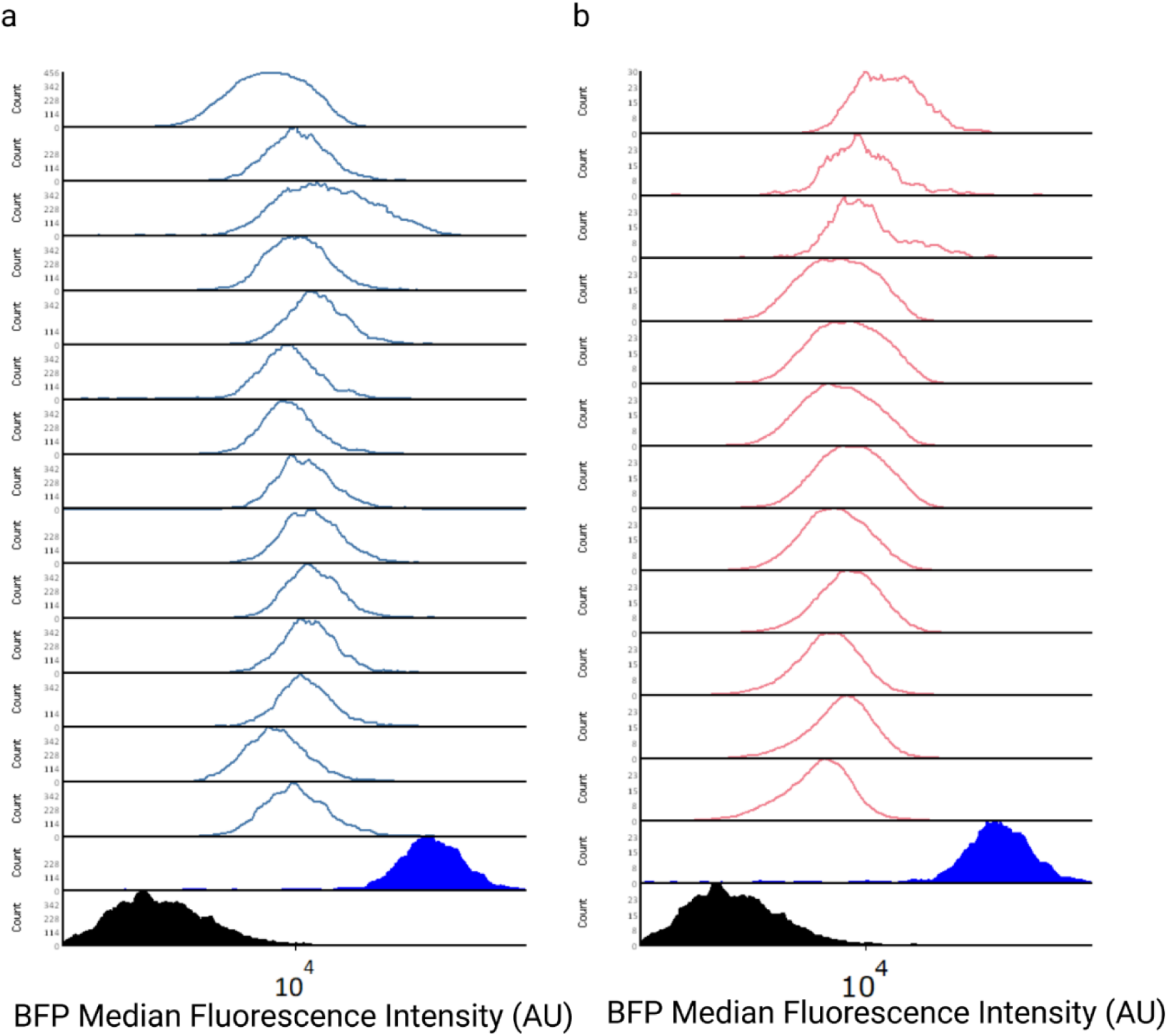
Median fluorescence intensity of cheater cultures was measured in a Beckman and Coulter Cytoflex with a 405 nm excitation laser and 450/45 bandpass to ensure growth was not due to contamination. **(a)** Histograms of strain expressing SpSwi6/SpClr4 (SGy162, blue lines) and **(b)** control strain (SGy139, red lines) show all strains are BFP positive after re-growth from experiments in Figure 1. CEN.PK2.1C wildtype control and BFP positive control filled in black and blue respectively.

**Extended Data Fig. 12.**
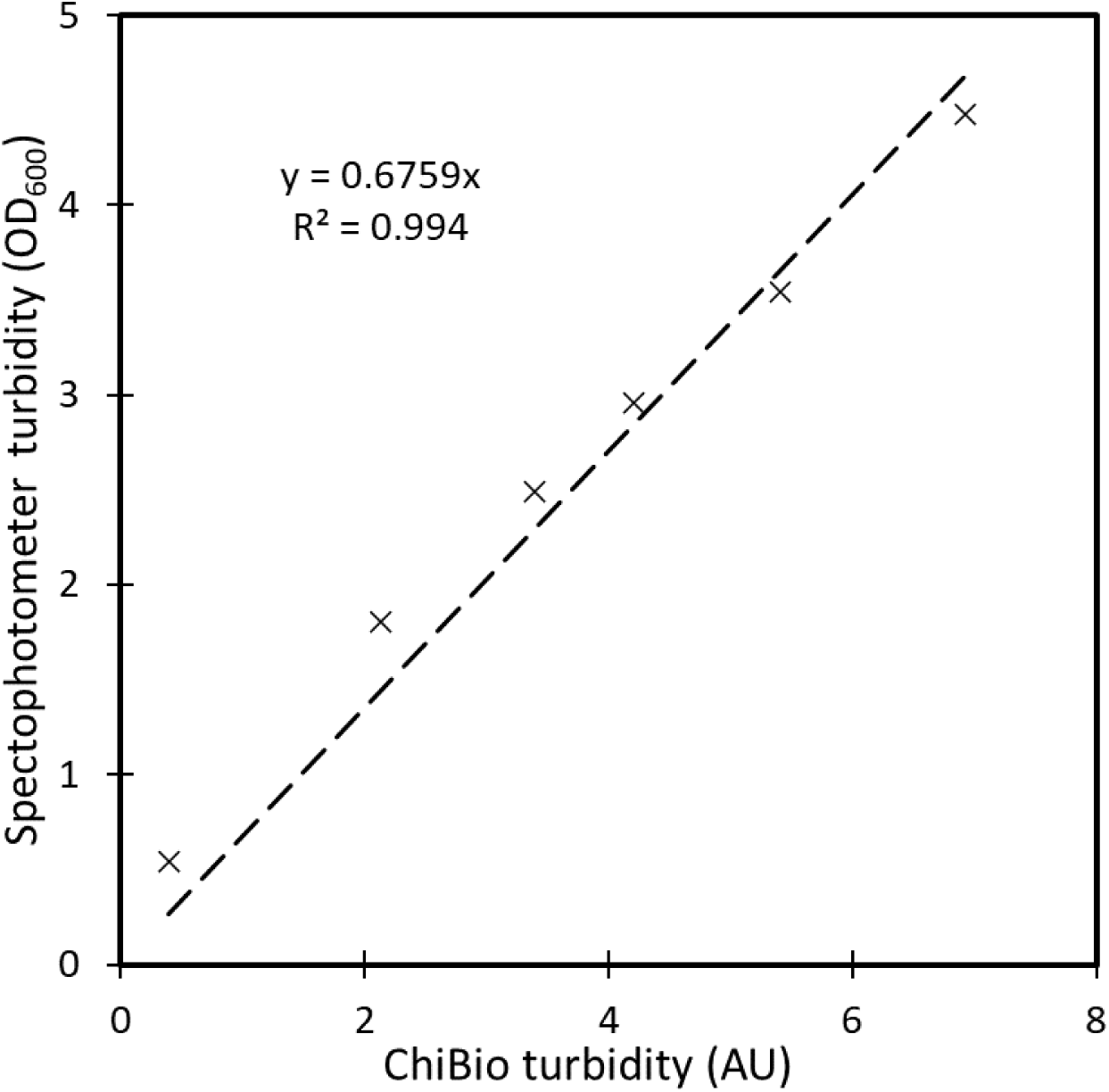
ChiBio OD calibration curve. CENPK2.1C strain was grown to saturation, serially diluted, and measured in both spectrophotometer and ChiBio. The data was plotted, and a linear regression was calculated, the linear equation was used to convert ChiBio turbidity to Spectrophotometer OD_600_ absorbance values.

**Extended Data Fig. 13.**
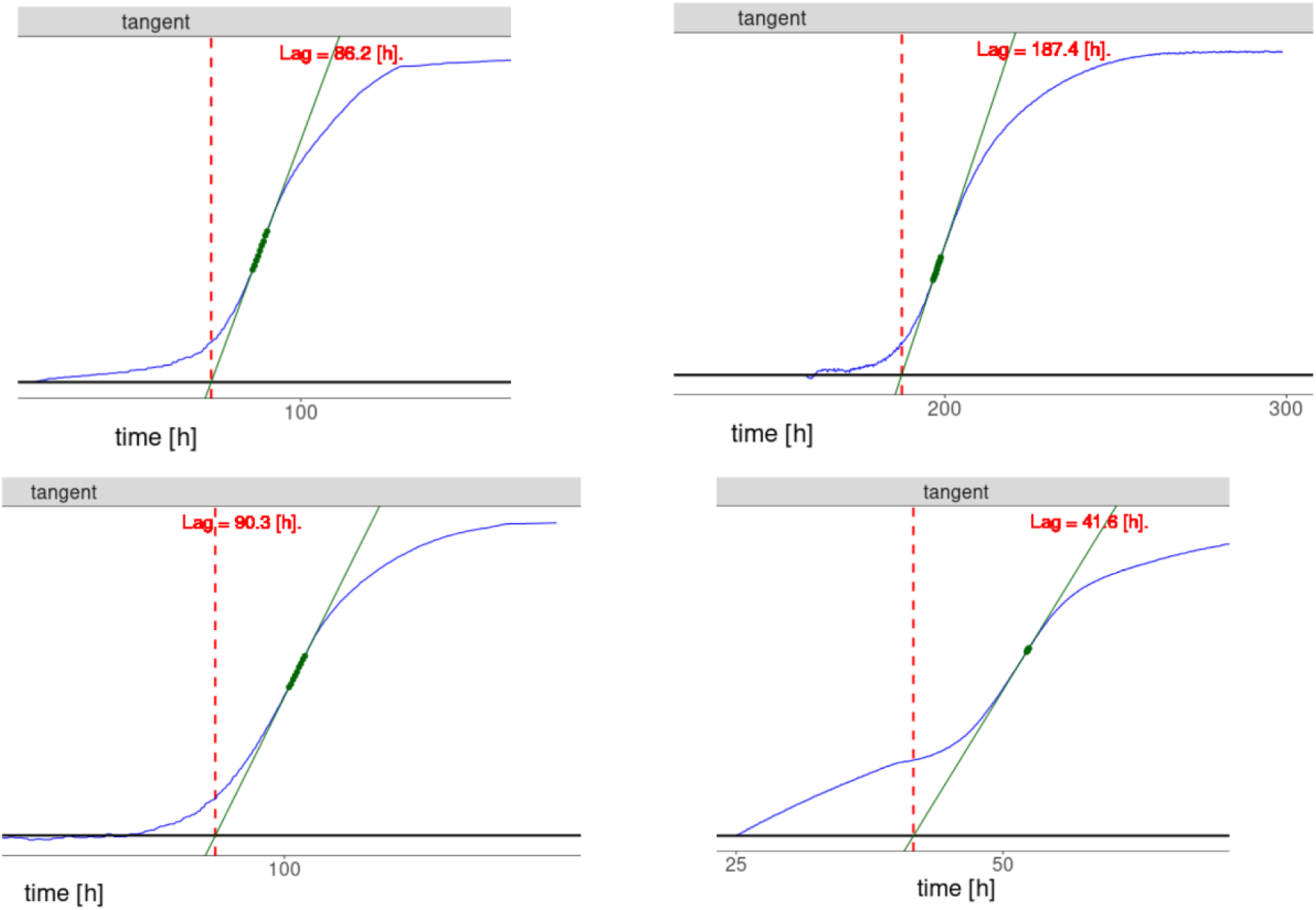
Representative plots used to calculate the growth lag phase of cheater cultures. Culture stability was estimated by measuring the duration of the lag phase using the tangent method^132^, in what should have been non-permissive conditions (darkness). To calculate the lag phase, we used the first observed maximum growth rate and a local regression of 9 points (green points) as parameters.

---

**Extended Data Table 1.**
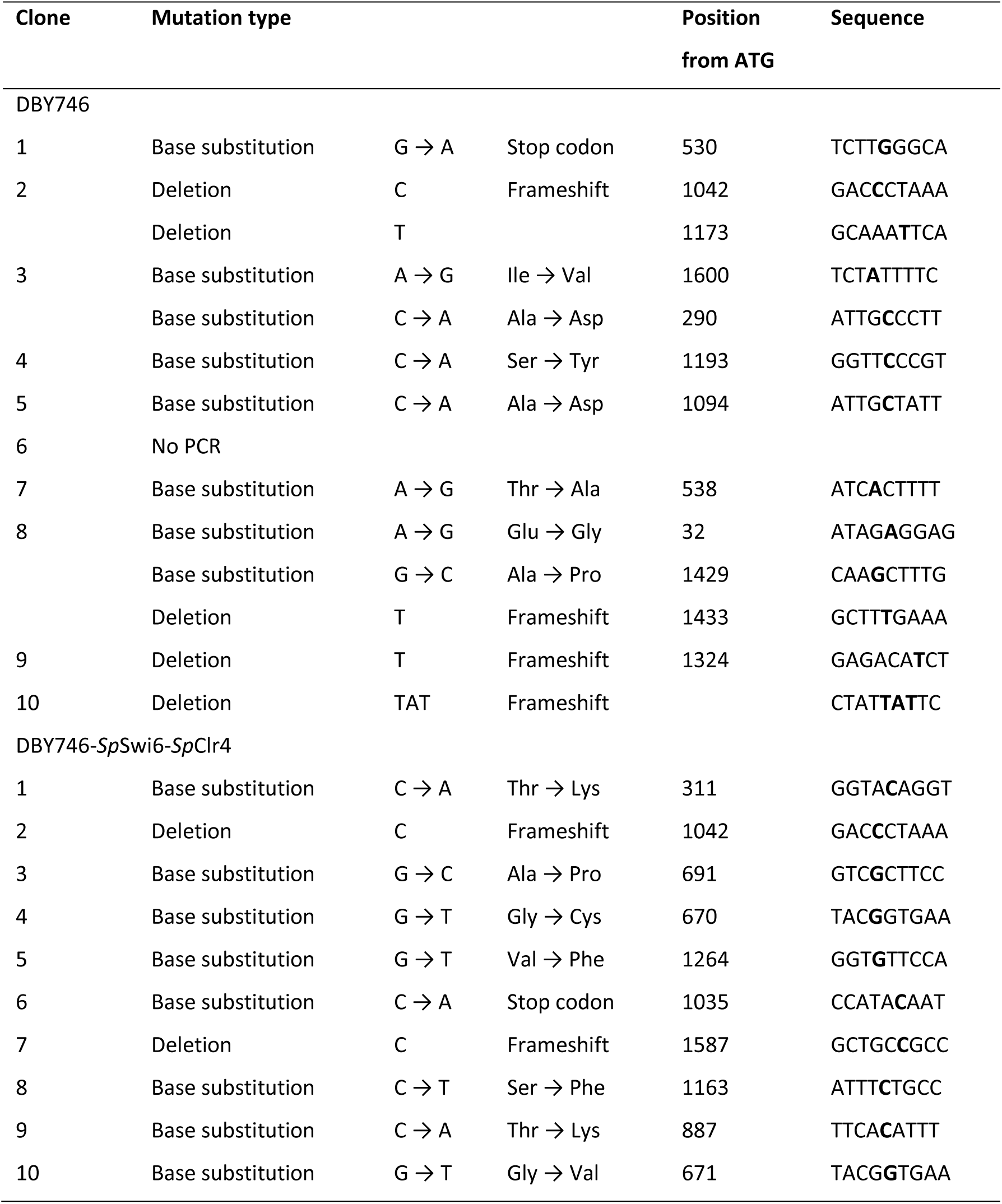
Mutation patterns obtained in Canr colonies from day 9 incubation.

**Extended Data Table 2.**
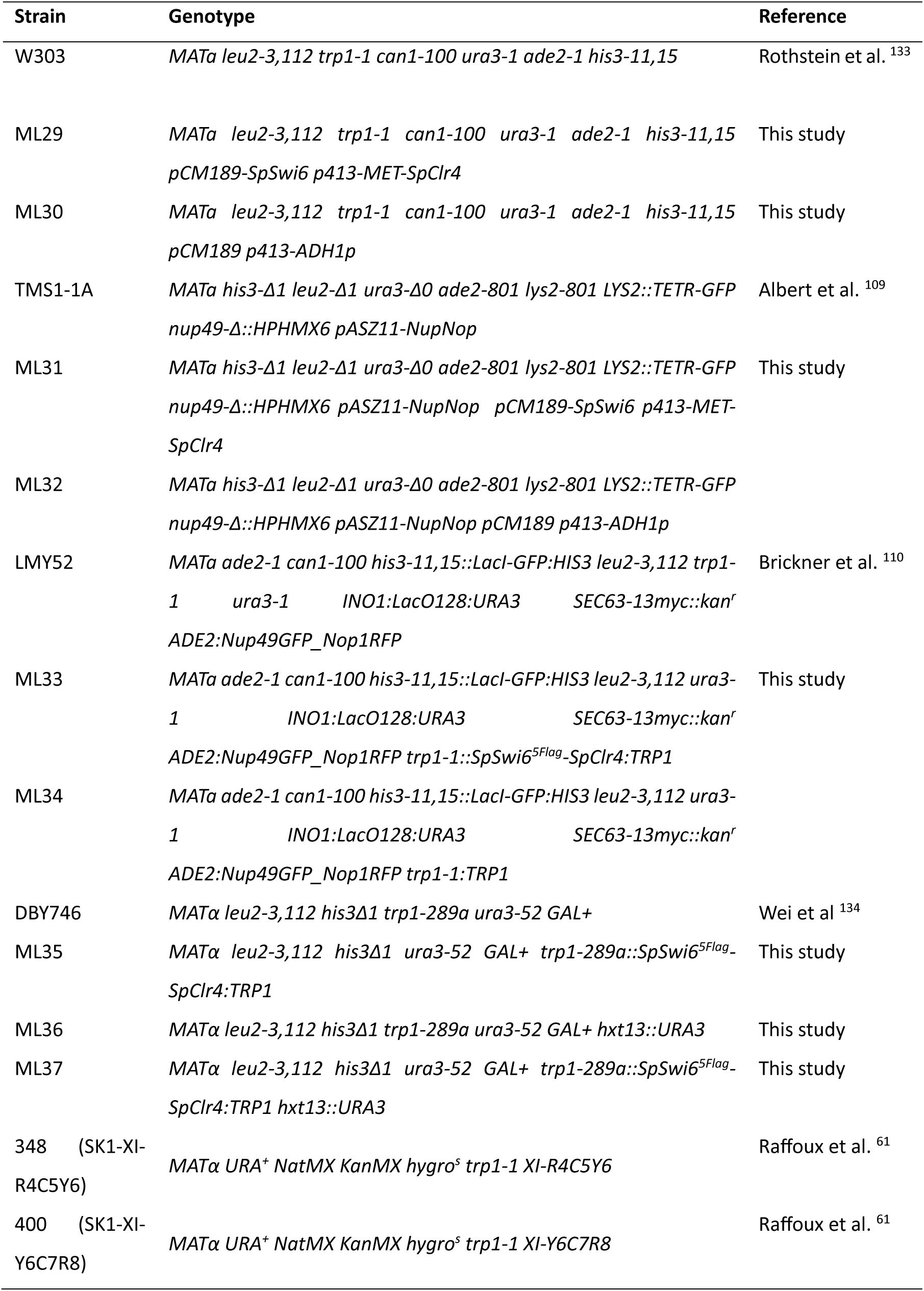
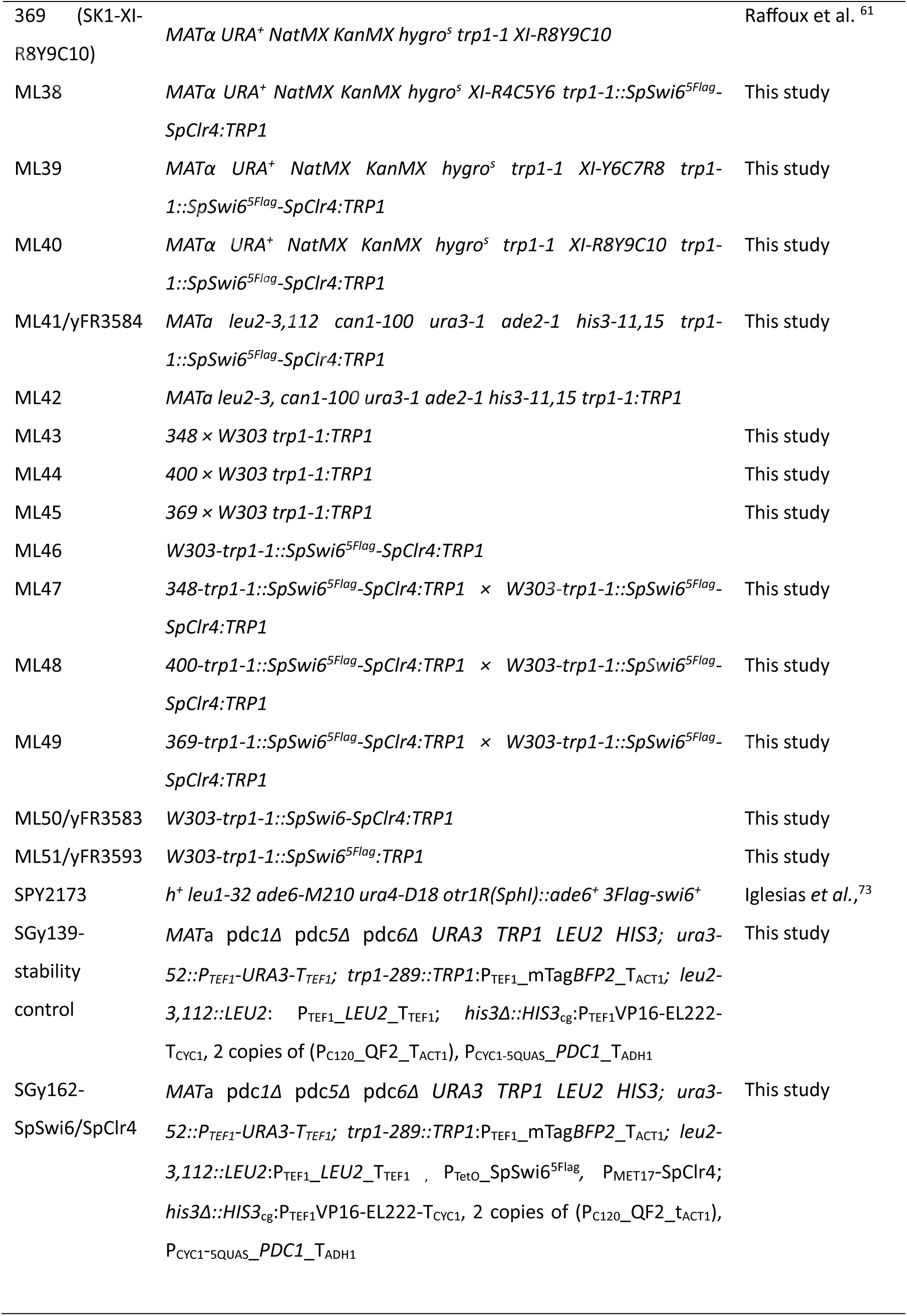
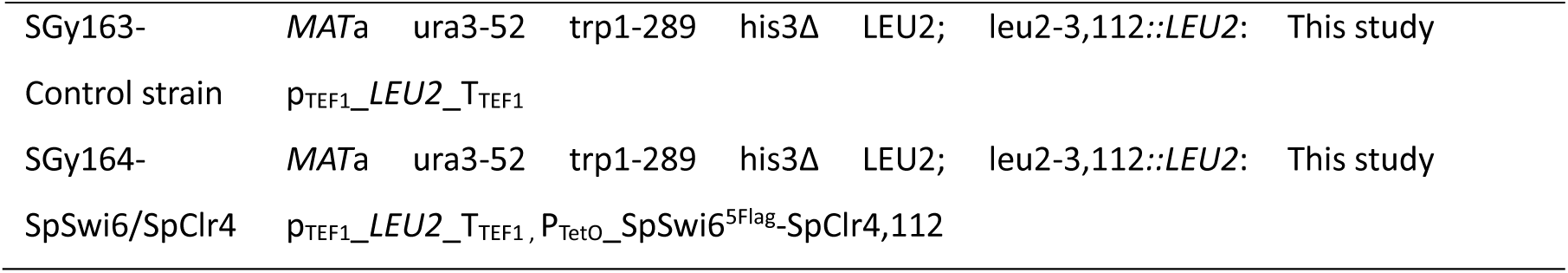
Yeast strains used in this study.

**Extended Data Table 3.**
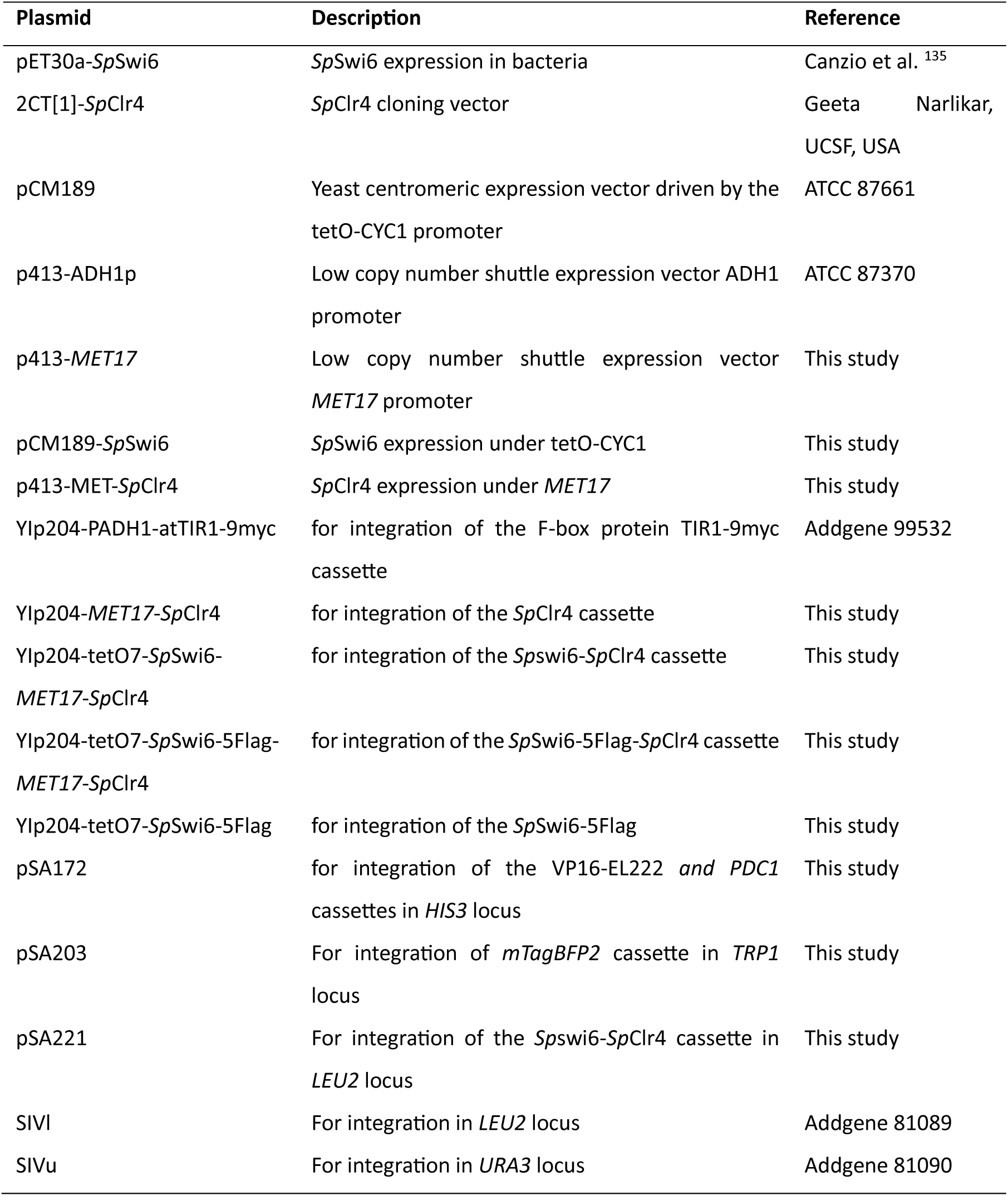
Plasmids used in this study.

## Supplementary Note 1

Many possible mutations could enable this strain to escape its light regulation, such as mutations on the light-dependent promoter (P_C120_) in the OptoAMP2 circuit (Fig. 6b) that make the promoter constitutive; translocation of the OptoAMP2 transcriptional activator (QF2) to a constitutive locus; *PDC1* gene amplification and/or translocation to a constitutive promoter; and less obvious mutations that break the engineered *PDC1* regulation. Previously reported mutations that allow triple *PDC* deletion strains to grow on glucose, such as mutations on *MTH1*^136^, are unlikely to be involved in the adaptation of our strains, as these mutations leave strains still dependent on a C2 carbon source for the cytosolic acetyl-CoA synthesis, which was not provided in our medium. Mutations leading to cheater strains may or may not have been preceded by deletion or mutation of *Sp*Swi6, *Sp*Clr4, or both. Identifying the nature and frequency of mutations present in each of our cheater strains, however, is beyond the scope of this study.

## Notes

### Competing Interest Statement

The authors have declared no competing interest.

## References

1 Schindler, D. Cell (2023). 10.1016/j.cell.2023.10.015

2 Taghon, G. J. & Strychalski, E. A. Cell Genom. (2023). 10.1016/j.xgen.2023.100438

3 Zhao, Y. Cell (2023). 10.1016/j.cell.2023.09.025

4 Cremer, T. & Cremer, C. Chromosome territories, nuclear architecture and gene regulation in mammalian cells. Nat Rev Genet 2, 292–301 (2001). 10.1038/35066075

5 Lieberman-Aiden, E. et al. Comprehensive mapping of long-range interactions reveals folding principles of the human genome. Science 326, 289–293 (2009). 10.1126/science.1181369

6 Dekker, J., Rippe, K., Dekker, M. & Kleckner, N. Capturing chromosome conformation. Science 295, 1306–1311 (2002). 10.1126/science.1067799

7 Duan, Z. et al. A three-dimensional model of the yeast genome. Nature 465, 363–367 (2010). 10.1038/nature08973

8 Sexton, T. et al. Three-dimensional folding and functional organization principles of the Drosophila genome. Cell 148, 458–472 (2012). 10.1016/j.cell.2012.01.010

9 Crane, E. et al. Condensin-driven remodelling of X chromosome topology during dosage compensation. Nature 523, 240–244 (2015). 10.1038/nature14450

10 Dong, Q. et al. Genome-wide Hi-C analysis reveals extensive hierarchical chromatin interactions in rice. Plant J 94, 1141–1156 (2018). 10.1111/tpj.13925

11 Dong, P. et al. 3D Chromatin Architecture of Large Plant Genomes Determined by Local A/B Compartments. Mol Plant 10, 1497–1509 (2017). 10.1016/j.molp.2017.11.005

12 Stephens, A. D. et al. Chromatin histone modifications and rigidity affect nuclear morphology independent of lamins. Mol Biol Cell 29, 220–233 (2018). 10.1091/mbc.E17-06-0410

13 Allshire, R. C. & Madhani, H. D. Ten principles of heterochromatin formation and function. Nat Rev Mol Cell Biol 19, 229–244 (2018). 10.1038/nrm.2017.119

14 Bannister, A. J. et al. Selective recognition of methylated lysine 9 on histone H3 by the HP1 chromo domain. Nature 410, 120–124 (2001). 10.1038/35065138

15 Nakayama, J., Rice, J. C., Strahl, B. D., Allis, C. D. & Grewal, S. I. Role of histone H3 lysine 9 methylation in epigenetic control of heterochromatin assembly. Science 292, 110–113 (2001). 10.1126/science.1060118

16 Stewart, M. D., Li, J. & Wong, J. Relationship between histone H3 lysine 9 methylation, transcription repression, and heterochromatin protein 1 recruitment. Mol Cell Biol 25, 2525–2538 (2005). 10.1128/mcb.25.7.2525-2538.2005

17 Sanulli, S. et al. HP1 reshapes nucleosome core to promote heterochromatin phase separation. Nature (2019). 10.1038/s41586-019-1669-2

18 Mizuguchi, T., Barrowman, J. & Grewal, S. I. S. Chromosome domain architecture and dynamic organization of the fission yeast genome. FEBS letters 589, 2975–2986 (2015). 10.1016/j.febslet.2015.06.008

19 Hickman, M. A., Froyd, C. A. & Rusche, L. N. Reinventing heterochromatin in budding yeasts: Sir2 and the origin recognition complex take center stage. Eukaryot Cell 10, 1183–1192 (2011). 10.1128/ec.05123-11

20 Lomberk, G., Wallrath, L. & Urrutia, R. The Heterochromatin Protein 1 family. Genome Biol 7, 228 (2006). 10.1186/gb-2006-7-7-228

21 Cai, S., Chen, C., Tan, Z. Y., Huang, Y., Shi, J. & Gan, L. Cryo-ET reveals the macromolecular reorganization of S. pombe mitotic chromosomes in vivo. Proceedings of the National Academy of Sciences of the United States of America 115, 10977–10982 (2018). 10.1073/pnas.1720476115

22 Chen, C. et al. Budding yeast chromatin is dispersed in a crowded nucleoplasm in vivo. Molecular biology of the cell 27, 3357–3368 (2016). 10.1091/mbc.E16-07-0506

23 Kakui, Y., Rabinowitz, A., Barry, D. J. & Uhlmann, F. Condensin-mediated remodeling of the mitotic chromatin landscape in fission yeast. Nature genetics 49, 1553–1557 (2017). 10.1038/ng.3938

24 Tanizawa, H., Kim, K. D., Iwasaki, O. & Noma, K. I. Architectural alterations of the fission yeast genome during the cell cycle. Nature structural & molecular biology 24, 965–976 (2017). 10.1038/nsmb.3482

25 Grunstein, M. & Gasser, S. M. Epigenetics in Saccharomyces cerevisiae. Cold Spring Harb Perspect Biol 5 (2013). 10.1101/cshperspect.a017491

26 Mizuguchi, T. et al. Cohesin-dependent globules and heterochromatin shape 3D genome architecture in S. pombe. Nature 516, 432–435 (2014). 10.1038/nature13833

27 Hsieh, T. H., Weiner, A., Lajoie, B., Dekker, J., Friedman, N. & Rando, O. J. Mapping Nucleosome Resolution Chromosome Folding in Yeast by Micro-C. Cell 162, 108–119 (2015). 10.1016/j.cell.2015.05.048

28 Linhoff, M. W., Garg, S. K. & Mandel, G. A high-resolution imaging approach to investigate chromatin architecture in complex tissues. Cell 163, 246–255 (2015). 10.1016/j.cell.2015.09.002

29 Imai, R. et al. Density imaging of heterochromatin in live cells using orientation-independent-DIC microscopy. Molecular biology of the cell 28, 3349–3359 (2017). 10.1091/mbc.E17-06-0359

30 Maison, C., Quivy, J. P., Probst, A. V. & Almouzni, G. Heterochromatin at mouse pericentromeres: a model for de novo heterochromatin formation and duplication during replication. Cold Spring Harbor symposia on quantitative biology 75, 155–165 (2010). 10.1101/sqb.2010.75.013

31 Saksouk, N., Simboeck, E. & Déjardin, J. Constitutive heterochromatin formation and transcription in mammals. Epigenetics & chromatin 8, 3 (2015). 10.1186/1756-8935-8-3

32 Bancaud, A., Huet, S., Daigle, N., Mozziconacci, J., Beaudouin, J. & Ellenberg, J. Molecular crowding affects diffusion and binding of nuclear proteins in heterochromatin and reveals the fractal organization of chromatin. The EMBO journal 28, 3785–3798 (2009). 10.1038/emboj.2009.340

33 Grewal, S. I. & Jia, S. Heterochromatin revisited. Nature reviews. Genetics 8, 35–46 (2007). 10.1038/nrg2008

34 Allis, C. D. & Jenuwein, T. The molecular hallmarks of epigenetic control. Nature reviews. Genetics 17, 487–500 (2016). 10.1038/nrg.2016.59

35 Brero, A. et al. Methyl CpG-binding proteins induce large-scale chromatin reorganization during terminal differentiation. The Journal of cell biology 169, 733–743 (2005). 10.1083/jcb.200502062

36 Nan, X., Tate, P., Li, E. & Bird, A. DNA methylation specifies chromosomal localization of MeCP2. Molecular and cellular biology 16, 414–421 (1996). 10.1128/mcb.16.1.414

37 Riley, R. et al. Comparative genomics of biotechnologically important yeasts. Proceedings of the National Academy of Sciences 113, 9882 (2016). 10.1073/pnas.1603941113

38 Spector, D. L. The dynamics of chromosome organization and gene regulation. Annu Rev Biochem 72, 573–608 (2003). 10.1146/annurev.biochem.72.121801.161724

39 Jin, Q. W., Fuchs, J. & Loidl, J. Centromere clustering is a major determinant of yeast interphase nuclear organization. J Cell Sci 113 **(Pt** **11**), 1903–1912 (2000).

40 Rabl, C. Uber Zelltheilung. Morphol. Jahrb. 10, 214–330 (1885).

41 Pouokam, M., Cruz, B., Burgess, S., Segal, M. R., Vazquez, M. & Arsuaga, J. The Rabl configuration limits topological entanglement of chromosomes in budding yeast. Scientific Reports 9, 6795 (2019). 10.1038/s41598-019-42967-4

42 Therizols, P., Duong, T., Dujon, B., Zimmer, C. & Fabre, E. Chromosome arm length and nuclear constraints determine the dynamic relationship of yeast subtelomeres. Proc Natl Acad Sci U S A 107, 2025–2030 (2010). 10.1073/pnas.0914187107

43 Berger, A. B. et al. High-resolution statistical mapping reveals gene territories in live yeast. Nat Methods 5, 1031–1037 (2008). 10.1038/nmeth.1266

44 Wong, H. et al. A predictive computational model of the dynamic 3D interphase yeast nucleus. Curr Biol 22, 1881–1890 (2012). 10.1016/j.cub.2012.07.069

45 Bystricky, K., Heun, P., Gehlen, L., Langowski, J. & Gasser, S. M. Long-range compaction and flexibility of interphase chromatin in budding yeast analyzed by high-resolution imaging techniques. Proc Natl Acad Sci U S A 101, 16495–16500 (2004). 10.1073/pnas.0402766101

46 Neumann, F. R. et al. Targeted INO80 enhances subnuclear chromatin movement and ectopic homologous recombination. Genes Dev 26, 369–383 (2012). 10.1101/gad.176156.111

47 Verdaasdonk, J. S. et al. Centromere tethering confines chromosome domains. Mol Cell 52, 819–831 (2013). 10.1016/j.molcel.2013.10.021

48 González, L. et al. Adaptive partitioning of a gene locus to the nuclear envelope in Saccharomyces cerevisiae is driven by polymer-polymer phase separation. Nature communications 14, 1135 (2023). 10.1038/s41467-023-36391-6

49 Zhu, Y. O., Siegal, M. L., Hall, D. W. & Petrov, D. A. Precise estimates of mutation rate and spectrum in yeast. Proc Natl Acad Sci U S A 111, E2310–2318 (2014). 10.1073/pnas.1323011111

50 Martincorena, I., Seshasayee, A. S. & Luscombe, N. M. Evidence of non-random mutation rates suggests an evolutionary risk management strategy. Nature 485, 95–98 (2012). 10.1038/nature10995

51 Measday, V. & Stirling, P. C. Navigating yeast genome maintenance with functional genomics. Briefings in Functional Genomics 15, 119–129 (2015). 10.1093/bfgp/elv033%J Briefings in Functional Genomics

52 Huang, M. E., Rio, A. G., Nicolas, A. & Kolodner, R. D. A genomewide screen in Saccharomyces cerevisiae for genes that suppress the accumulation of mutations. Proc Natl Acad Sci U S A 100, 11529–11534 (2003). 10.1073/pnas.2035018100

53 Lang, G. I. & Murray, A. W. Estimating the per-base-pair mutation rate in the yeast Saccharomyces cerevisiae. Genetics 178, 67–82 (2008). 10.1534/genetics.107.071506

54 Stirling, P. C., Shen, Y., Corbett, R., Jones, S. J. M. & Hieter, P. Genome destabilizing mutator alleles drive specific mutational trajectories in Saccharomyces cerevisiae. Genetics 196, 403–412 (2014). 10.1534/genetics.113.159806

55 Fabrizio, P. et al. Superoxide is a mediator of an altruistic aging program in Saccharomyces cerevisiae. J Cell Biol 166, 1055–1067 (2004). 10.1083/jcb.200404002

56 Smith, S., Hwang, J. Y., Banerjee, S., Majeed, A., Gupta, A. & Myung, K. Mutator genes for suppression of gross chromosomal rearrangements identified by a genome-wide screening in Saccharomyces cerevisiae. Proc Natl Acad Sci U S A 101, 9039–9044 (2004). 10.1073/pnas.0403093101

57 Kanellis, P. et al. A screen for suppressors of gross chromosomal rearrangements identifies a conserved role for PLP in preventing DNA lesions. PLoS Genet 3, e134 (2007). 10.1371/journal.pgen.0030134

58 Madia, F. et al. Longevity mutation in SCH9 prevents recombination errors and premature genomic instability in a Werner/Bloom model system. The Journal of cell biology 180, 67–81 (2008). 10.1083/jcb.200707154

59 Chen, C. & Kolodner, R. D. Gross chromosomal rearrangements in Saccharomyces cerevisiae replication and recombination defective mutants. Nature genetics 23, 81–85 (1999). 10.1038/12687

60 Putnam, C. D. & Kolodner, R. D. Determination of gross chromosomal rearrangement rates. Cold Spring Harbor protocols 2010, pdb.prot5492 (2010). 10.1101/pdb.prot5492

61 Raffoux, X., Bourge, M., Dumas, F., Martin, O. C. & Falque, M. High-throughput measurement of recombination rates and genetic interference in Saccharomyces cerevisiae. Yeast 35, 431–442 (2018). 10.1002/yea.3315

62 Lichten, M. & Haber, J. E. Position effects in ectopic and allelic mitotic recombination in Saccharomyces cerevisiae. Genetics 123, 261–268 (1989). 10.1093/genetics/123.2.261

63 Datta, A., Adjiri, A., New, L., Crouse, G. F. & Jinks Robertson, S. Mitotic crossovers between diverged sequences are regulated by mismatch repair proteins in Saccaromyces cerevisiae. Mol Cell Biol 16, 1085–1093 (1996). 10.1128/MCB.16.3.1085

64 McIlwraith, M. J. & West, S. C. DNA Repair Synthesis Facilitates RAD52-Mediated Second-End Capture during DSB Repair. Molecular Cell 29, 510–516 (2008). 10.1016/j.molcel.2007.11.037

65 She, R. & Jarosz, D. F. Mapping Causal Variants with Single-Nucleotide Resolution Reveals Biochemical Drivers of Phenotypic Change. Cell 172, 478–490.e415 (2018). 10.1016/j.cell.2017.12.015

66 Ji, Q., Mai, J., Ding, Y., Wei, Y., Ledesma-Amaro, R. & Ji, X.-J. Improving the homologous recombination efficiency of Yarrowia lipolytica by grafting heterologous component from Saccharomyces cerevisiae. Metabolic engineering communications 11, e00152–e00152 (2020). 10.1016/j.mec.2020.e00152

67 Cromie, G. & Smith, G. R. Meiotic Recombination in Schizosaccharomyces pombe: A Paradigm for Genetic and Molecular Analysis. Genome Dyn Stab 3, 195 (2008). 10.1007/7050_2007_025

68 Virgin, J. B. & Bailey, J. P. The M26 hotspot of Schizosaccharomyces pombe stimulates meiotic ectopic recombination and chromosomal rearrangements. Genetics 149, 1191–1204 (1998). 10.1093/genetics/149.3.1191

69 Wilfert, L., Gadau, J. & Schmid-Hempel, P. Variation in genomic recombination rates among animal taxa and the case of social insects. Heredity (Edinb) 98, 189–197 (2007). 10.1038/sj.hdy.6800950

70 Segura, J. et al. Evolution of recombination in eutherian mammals: insights into mechanisms that affect recombination rates and crossover interference. Proc Biol Sci 280, 20131945 (2013). 10.1098/rspb.2013.1945

71 Tiley, G. P. & Burleigh, J. G. The relationship of recombination rate, genome structure, and patterns of molecular evolution across angiosperms. BMC Evol Biol 15, 194 (2015). 10.1186/s12862-015-0473-3

72 Barton, A. B., Pekosz, M. R., Kurvathi, R. S. & Kaback, D. B. Meiotic recombination at the ends of chromosomes in Saccharomyces cerevisiae. Genetics 179, 1221–1235 (2008). 10.1534/genetics.107.083493

73 Iglesias, N. et al. Native Chromatin Proteomics Reveals a Role for Specific Nucleoporins in Heterochromatin Organization and Maintenance. Molecular cell 77, 51–66.e58 (2020). 10.1016/j.molcel.2019.10.018

74 Grewal, S. I. S. The molecular basis of heterochromatin assembly and epigenetic inheritance. Molecular cell 83, 1767–1785 (2023). 10.1016/j.molcel.2023.04.020

75 Jeronimo, C. & Robert, F. Kin28 regulates the transient association of Mediator with core promoters. Nature structural & molecular biology 21, 449–455 (2014). 10.1038/nsmb.2810

76 Park, D., Lee, Y., Bhupindersingh, G. & Iyer, V. R. Widespread misinterpretable ChIP-seq bias in yeast. PloS one 8, e83506 (2013). 10.1371/journal.pone.0083506

77 Teytelman, L., Thurtle, D. M., Rine, J. & van Oudenaarden, A. Highly expressed loci are vulnerable to misleading ChIP localization of multiple unrelated proteins. Proceedings of the National Academy of Sciences of the United States of America 110, 18602–18607 (2013). 10.1073/pnas.1316064110

78 Jih, G. et al. Unique roles for histone H3K9me states in RNAi and heritable silencing of transcription. Nature 547, 463–467 (2017). 10.1038/nature23267

79 Kuzdere, T., Flury, V., Schalch, T., Iesmantavicius, V., Hess, D. & Bühler, M. Differential phosphorylation of Clr4(SUV39H) by Cdk1 accompanies a histone H3 methylation switch that is essential for gametogenesis. EMBO reports 24, e55928 (2023). 10.15252/embr.202255928

80 Lalwani, M. A., Zhao, E. M. & Avalos, J. L. Current and future modalities of dynamic control in metabolic engineering. Curr Opin Biotechnol 52, 56–65 (2018). 10.1016/j.copbio.2018.02.007

81 Zhao, E. M. et al. Optogenetic regulation of engineered cellular metabolism for microbial chemical production. Nature 555, 683–687 (2018). 10.1038/nature26141

82 Lalwani, M. A., Zhao, E. M., Wegner, S. A. & Avalos, J. L. The Neurospora crassa Inducible Q System Enables Simultaneous Optogenetic Amplification and Inversion in Saccharomyces cerevisiae for Bidirectional Control of Gene Expression. ACS synthetic biology 10, 2060–2075 (2021). 10.1021/acssynbio.1c00229

83 Steel, H., Habgood, R., Kelly, C. L. & Papachristodoulou, A. In situ characterisation and manipulation of biological systems with Chi.Bio. PLoS biology 18, e3000794 (2020). 10.1371/journal.pbio.3000794

84 Nash, A. I. et al. Structural basis of photosensitivity in a bacterial light-oxygen-voltage/helix-turn-helix (LOV-HTH) DNA-binding protein. Proceedings of the National Academy of Sciences of the United States of America 108, 9449–9454 (2011). 10.1073/pnas.1100262108

85 Makova, K. D. & Hardison, R. C. The effects of chromatin organization on variation in mutation rates in the genome. Nature reviews. Genetics 16, 213–223 (2015). 10.1038/nrg3890

86 Pich, O., Muiños, F., Sabarinathan, R., Reyes-Salazar, I., Gonzalez-Perez, A. & Lopez-Bigas, N. Somatic and Germline Mutation Periodicity Follow the Orientation of the DNA Minor Groove around Nucleosomes. Cell 175, 1074–1087.e1018 (2018). 10.1016/j.cell.2018.10.004

87 Montavon, T. et al. Complete loss of H3K9 methylation dissolves mouse heterochromatin organization. Nature communications 12, 4359 (2021). 10.1038/s41467-021-24532-8

88 Henderson, I. R. & Bomblies, K. Evolution and Plasticity of Genome-Wide Meiotic Recombination Rates. Annu Rev Genet 55, 23–43 (2021). 10.1146/annurev-genet-021721-033821

89 Nicklas, R. B. Chance encounters and precision in mitosis. J Cell Sci 89 **(Pt** **3**), 283–285 (1988). 10.1242/jcs.89.3.283

90 Goloborodko, A., Marko, J. F. & Mirny, L. A. Chromosome Compaction by Active Loop Extrusion. Biophysical journal 110, 2162–2168 (2016). 10.1016/j.bpj.2016.02.041

91 Ganji, M. et al. Real-time imaging of DNA loop extrusion by condensin. Science (New York, N.Y.) 360, 102–105 (2018). 10.1126/science.aar7831

92 Davidson, I. F., Bauer, B., Goetz, D., Tang, W., Wutz, G. & Peters, J. M. DNA loop extrusion by human cohesin. Science (New York, N.Y.) 366, 1338–1345 (2019). 10.1126/science.aaz3418

93 Kim, Y., Shi, Z., Zhang, H., Finkelstein, I. J. & Yu, H. Human cohesin compacts DNA by loop extrusion. *Science (New York*, N.Y.) 366, 1345–1349 (2019). 10.1126/science.aaz4475

94 Murugan, A., Huse, D. A. & Leibler, S. Speed, dissipation, and error in kinetic proofreading. Proc Natl Acad Sci U S A 109, 12034–12039 (2012). 10.1073/pnas.1119911109

95 Tlusty, T., Bar-Ziv, R. & Libchaber, A. High-fidelity DNA sensing by protein binding fluctuations. Phys Rev Lett 93, 258103 (2004). 10.1103/PhysRevLett.93.258103

96 Cox, M. M. Regulation of bacterial RecA protein function. Crit Rev Biochem Mol Biol 42, 41–63 (2007). 10.1080/10409230701260258

97 Lenski, R. E., Rose, M. R., Simpson, S. C. & Tadler, S. C. Long-Term Experimental Evolution in Escherichia coli. I. Adaptation and Divergence During 2,000 Generations. The American Naturalist 138, 1315–1341 (1991).

98 Blount, Z. D., Barrick, J. E., Davidson, C. J. & Lenski, R. E. Genomic analysis of a key innovation in an experimental population. Nature 489, 513-+ (2012). 10.1038/nature11514

99 Yona, A. H. et al. Chromosomal duplication is a transient evolutionary solution to stress. Proceedings of the National Academy of Sciences of the United States of America 109, 21010–21015 (2012). 10.1073/pnas.1211150109

100 Matic, I. Mutation Rate Heterogeneity Increases Odds of Survival in Unpredictable Environments. Molecular cell 75, 421–425 (2019). 10.1016/j.molcel.2019.06.029

101 Basso, L. C., de Amorim, H. V., de Oliveira, A. J. & Lopes, M. L. Yeast selection for fuel ethanol production in Brazil. FEMS Yeast Res 8, 1155–1163 (2008). 10.1111/j.1567-1364.2008.00428.x

102 Carro, D. & Piña, B. Genetic analysis of the karyotype instability in natural wine yeast strains. *Yeast (Chichester*, England) 18, 1457–1470 (2001). 10.1002/yea.789

103 Querol, A. & Bond, U. The complex and dynamic genomes of industrial yeasts. FEMS Microbiol Lett 293, 1–10 (2009). 10.1111/j.1574-6968.2008.01480.x

104 Rodrigues-Prause, A. et al. A Case Study of Genomic Instability in an Industrial Strain of Saccharomyces cerevisiae. G3 (Bethesda) 8, 3703–3713 (2018). 10.1534/g3.118.200446

105 Argueso, J. L. et al. Genome structure of a Saccharomyces cerevisiae strain widely used in bioethanol production. Genome Res 19, 2258–2270 (2009). 10.1101/gr.091777.109

106 Borneman, A. R. et al. Whole-genome comparison reveals novel genetic elements that characterize the genome of industrial strains of Saccharomyces cerevisiae. PLoS Genet 7, e1001287 (2011). 10.1371/journal.pgen.1001287

107 Steensels, J., Snoek, T., Meersman, E., Picca Nicolino, M., Voordeckers, K. & Verstrepen, K. J. Improving industrial yeast strains: exploiting natural and artificial diversity. FEMS Microbiol Rev 38, 947–995 (2014). 10.1111/1574-6976.12073

108 Vaishnav, E. D. et al. The evolution, evolvability and engineering of gene regulatory DNA. Nature 603, 455–463 (2022). 10.1038/s41586-022-04506-6

109 Albert, B. et al. Systematic characterization of the conformation and dynamics of budding yeast chromosome XII. The Journal of cell biology 202, 201–210 (2013). 10.1083/jcb.201208186

110 Brickner, D. G. et al. Transcription factor binding to a DNA zip code controls interchromosomal clustering at the nuclear periphery. Developmental cell 22, 1234–1246 (2012). 10.1016/j.devcel.2012.03.012

111 Meilhoc, E. & Teissie, J. Electrotransformation of Saccharomyces cerevisiae. *Methods in molecular biology (Clifton*, N.J.) 2050, 187–193 (2020). 10.1007/978-1-4939-9740-4_21

112 Schindelin, J., et al. Fiji: an open-source platform for biological-image analysis. Nature methods 9, 676–682 (2012). 10.1038/nmeth.2019

113 Kechkar, A., Nair, D., Heilemann, M., Choquet, D. & Sibarita, J. B. Real-time analysis and visualization for single-molecule based super-resolution microscopy. PloS one 8, e62918 (2013). 10.1371/journal.pone.0062918

114 Verdaasdonk, J. S., Gardner, R., Stephens, A. D., Yeh, E. & Bloom, K. Tension-dependent nucleosome remodeling at the pericentromere in yeast. Molecular biology of the cell 23, 2560–2570 (2012). 10.1091/mbc.E11-07-0651

115 Scheffold, F., Diaz-Leyva, P., Reufer, M., Ben Braham, N., Lynch, I. & Harden, J. L. Brushlike interactions between thermoresponsive microgel particles. Phys Rev Lett 104, 128304 (2010). 10.1103/PhysRevLett.104.128304

116 Lõoke, M., Kristjuhan, K. & Kristjuhan, A. Extraction of genomic DNA from yeasts for PCR-based applications. BioTechniques 50, 325–328 (2011). 10.2144/000113672

117 Langmead, B. & Salzberg, S. L. Fast gapped-read alignment with Bowtie 2. Nature Methods 9, 357–U354 (2012). 10.1038/Nmeth.1923

118 Li, H. et al. The Sequence Alignment/Map format and SAMtools. Bioinformatics 25, 2078–2079 (2009). 10.1093/bioinformatics/btp352

119 Danecek, P. et al. The variant call format and VCFtools. Bioinformatics 27, 2156–2158 (2011). 10.1093/bioinformatics/btr330

120 Li, H. A statistical framework for SNP calling, mutation discovery, association mapping and population genetical parameter estimation from sequencing data. Bioinformatics 27, 2987–2993 (2011). 10.1093/bioinformatics/btr509

121 Jeronimo, C. et al. FACT is recruited to the +1 nucleosome of transcribed genes and spreads in a Chd1-dependent manner. Molecular cell 81, 3542–3559.e3511 (2021). 10.1016/j.molcel.2021.07.010

122 Bolger, A. M., Lohse, M. & Usadel, B. Trimmomatic: a flexible trimmer for Illumina sequence data. Bioinformatics 30, 2114–2120 (2014). 10.1093/bioinformatics/btu170

123 Langmead, B., Trapnell, C., Pop, M. & Salzberg, S. L. Ultrafast and memory-efficient alignment of short DNA sequences to the human genome. Genome biology 10, R25 (2009). 10.1186/gb-2009-10-3-r25

124 Li, H. et al. The Sequence Alignment/Map format and SAMtools. Bioinformatics 25, 2078–2079 (2009). 10.1093/bioinformatics/btp352

125 Quinlan, A. R. & Hall, I. M. BEDTools: a flexible suite of utilities for comparing genomic features. Bioinformatics 26, 841–842 (2010). 10.1093/bioinformatics/btq033

126 Casper, J. et al. The UCSC Genome Browser database: 2018 update. Nucleic acids research 46, D762–d769 (2018). 10.1093/nar/gkx1020

127 Ramírez, F. et al. deepTools2: a next generation web server for deep-sequencing data analysis. Nucleic acids research 44, W160–165 (2016). 10.1093/nar/gkw257

128 Brunelle, M. et al. Aggregate and Heatmap Representations of Genome-Wide Localization Data Using VAP, a Versatile Aggregate Profiler. *Methods in molecular biology (Clifton*, N.J.) 1334, 273–298 (2015). 10.1007/978-1-4939-2877-4_18

129 Coulombe, C. et al. VAP: a versatile aggregate profiler for efficient genome-wide data representation and discovery. Nucleic acids research 42, W485–493 (2014). 10.1093/nar/gku302

130 Pelechano, V., Wei, W. & Steinmetz, L. M. Extensive transcriptional heterogeneity revealed by isoform profiling. Nature 497, 127–131 (2013). 10.1038/nature12121

131 Gresham, D. et al. The repertoire and dynamics of evolutionary adaptations to controlled nutrient-limited environments in yeast. PLoS Genet 4, e1000303 (2008). 10.1371/journal.pgen.1000303

132 Opalek, M., Smug, B. J. & Wloch-Salamon, D. How to determine microbial lag phase duration?, 2022.2011.2016.516631 (2022). 10.1101/2022.11.16.516631 %J bioRxiv

133 Rothstein, R. J. One-step gene disruption in yeast. Methods in enzymology 101, 202–211 (1983). 10.1016/0076-6879(83)01015-0

134 Wei, M. et al. Life span extension by calorie restriction depends on Rim15 and transcription factors downstream of Ras/PKA, Tor, and Sch9. PLoS genetics 4, e13–e13 (2008). 10.1371/journal.pgen.0040013

135 Canzio, D. et al. Chromodomain-mediated oligomerization of HP1 suggests a nucleosome-bridging mechanism for heterochromatin assembly. Molecular cell 41, 67–81 (2011). 10.1016/j.molcel.2010.12.016

136 Oud, B. et al. An internal deletion in MTH1 enables growth on glucose of pyruvate-decarboxylase negative, non-fermentative Saccharomyces cerevisiae. Microb Cell Fact 11, 131 (2012). 10.1186/1475-2859-11-131

